# Dynamic multi-omics and mechanistic modeling approach uncovers novel mechanisms of kidney fibrosis progression

**DOI:** 10.1101/2024.10.15.618507

**Authors:** Nadine Tuechler, Mira Lea Burtscher, Martin Garrido-Rodriguez, Muzamil Majid Khan, Denes Türei, Christian Tischer, Sarah Kaspar, Jennifer Jasmin Schwarz, Frank Stein, Mandy Rettel, Rafael Kramann, Mikhail M Savitski, Julio Saez-Rodriguez, Rainer Pepperkok

## Abstract

Kidney fibrosis, characterized by excessive extracellular matrix deposition, is a progressive disease that, despite affecting 10% of the population, lacks specific treatments and suitable biomarkers. This study presents a comprehensive, time-resolved multi-omics analysis of kidney fibrosis using an *in vitro* model system based on human kidney PDGFRβ^+^ mesenchymal cells aimed at unraveling disease mechanisms. Using transcriptomics, proteomics, phosphoproteomics, and secretomics we quantified over 14,000 biomolecules across seven time points following TGF-β stimulation. This revealed distinct temporal patterns in the expression and activity of known and potential kidney fibrosis markers and modulators. Data integration resulted in time-resolved multi-omic network models which allowed us to propose mechanisms related to fibrosis progression through early transcriptional reprogramming. Using siRNA knockdowns and phenotypic assays, we validated predictions and regulatory mechanisms underlying kidney fibrosis. In particular, we show that several early-activated transcription factors, including FLI1 and E2F1, act as negative regulators of collagen deposition and propose underlying molecular mechanisms. This work advances our understanding of the pathogenesis of kidney fibrosis and provides a resource to be further leveraged by the community.

Graphical Abstract

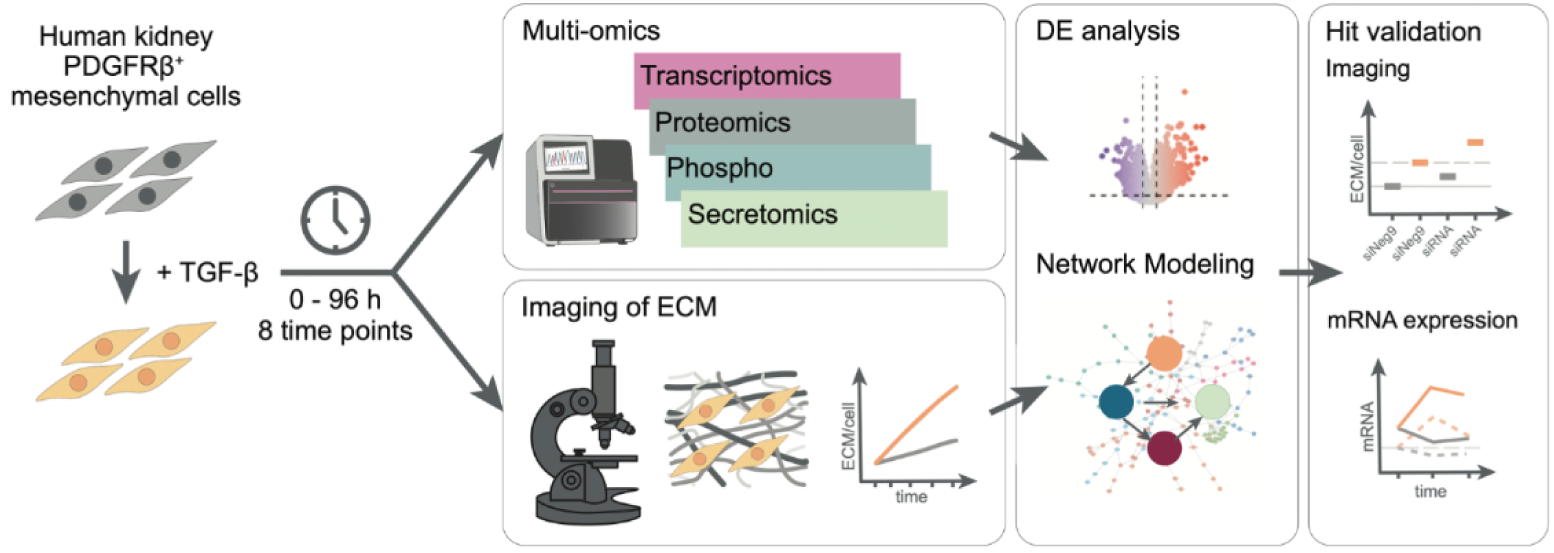

## 1. Introduction

Kidney fibrosis is a characteristic component of chronic kidney disease (CKD), characterized by the progressive accumulation and remodeling of extracellular matrix (ECM) components (Kim et al. 2022; Yamashita & Kramann 2024). This pathogenic process ultimately leads to disruption of tissue architecture and organ failure (Friedman et al. 2013; Kramann & Humphreys 2014; Mullins et al. 2016). Despite the high prevalence and severity of CKD, there are no specific treatments available and the few existing biomarkers are either impractical or too expensive for routine use, such as CKD273 (Argilés et al. 2013; Cañadas-Garre et al. 2018; Huang et al. 2023; Lousa et al. 2020; Prakash & Pinzani 2017; Rasmussen et al. 2019; Rodríguez-Ortiz et al. 2018; Verbeke et al. 2021; Yamashita & Kramann 2024; Yan et al. 2021). This gap in clinical management of the disease underscores the urgent need for a deeper understanding of the underlying molecular mechanisms.

Central to the pathogenesis of kidney fibrosis is the transdifferentiation of various cell types, including tissue-resident pericytes and fibroblasts, into myofibroblasts (Kendall & Feghali-Bostwick 2014). Activated myofibroblasts serve as the primary source of ECM components, driving the fibrotic process (Kim et al. 2022; Kramann & Humphreys 2014; Kuppe et al. 2021). Recent studies have shed light on cellular plasticity during fibrosis progression, but many aspects of the complexity of the disease remain poorly understood due to limitations such as single readouts, single time points, the inherent limitations of mouse models and the limited availability of human tissue samples (Liang & Liu 2023; Prakash & Pinzani 2017; Wei et al. 2024), (Kuppe et al. 2021).

Transcription factors (TFs) play a crucial role in myofibroblast differentiation and activation but remain challenging targets for drug development (Edeling et al. 2016; Humphreys 2018; Zhang et al. 2018); (Chang et al. 2012; Edeling et al. 2016; Humphreys 2018; Yamashita & Kramann 2024; Zhang et al. 2018). This highlights the importance of exploring novel approaches to modulate TF activity as well as upstream signaling processes and downstream phenotypic effects in the context of fibrosis.

Omics technologies combined with advanced computational methods offer powerful tools for studying complex processes like fibrosis progression (Lindenmeyer et al. 2021; Niewczas et al. 2019; Saez-Rodriguez et al. 2019). These include transcriptomics for tracking cellular differentiation, proteomics and phosphoproteomics for insights into signaling pathways, and secretomics for measuring related ECM and signaling components. Incorporating temporal resolution and integrating with phenotypic readouts are crucial for linking these molecular mechanisms to cellular and tissue-level processes.

In this study, we use transforming growth factor beta (TGF-β) to induce fibrotic progression in PDGFRβ+ human kidney mesenchymal cells and investigate the underlying molecular mechanisms in detail. This addresses many of the limitations of existing approaches by providing i) a perturbable system, ii) a robust phenotypic readout of ECM deposition and collagen secretion, iii) comprehensive multi-omics profiling of TGF-β-induced myofibroblast differentiation with high temporal resolution, and iv) a mechanistic, multi-omics network-based integration approach to generate testable hypotheses.

By leveraging this *in vitro* model system and our integrative analysis approach, we uncovered new mechanistic insights into the pathogenesis of kidney fibrosis and identified potential biomarkers and therapeutic targets for this challenging disease.

## 2. Results

### 2.1 Phenotypic and molecular features defining fibrosis in PDGFRβ^+^ mesenchymal cells over time

In this study, we present a comprehensive investigation of kidney fibrosis using an *in vitro* model system that enables detailed phenotypic and molecular disease characterization and has previously been used in the context of kidney fibrosis research (Kuppe et al. 2021). The use of human PDGFRβ^+^ mesenchymal cells in this tissue culture setup enables the induction of fibrotic differentiation with accelerated collagen deposition and ECM remodeling after stimulation with TGF-β (Chen et al. 2009; Coentro et al. 2021; Khan et al. 2023; Rønnow et al. 2020). This system not only enables the time-resolved characterization of the phenotypic and molecular features of fibrotic processes, but also facilitates subsequent perturbation and validation (Figure 1A).

**Figure 1:**
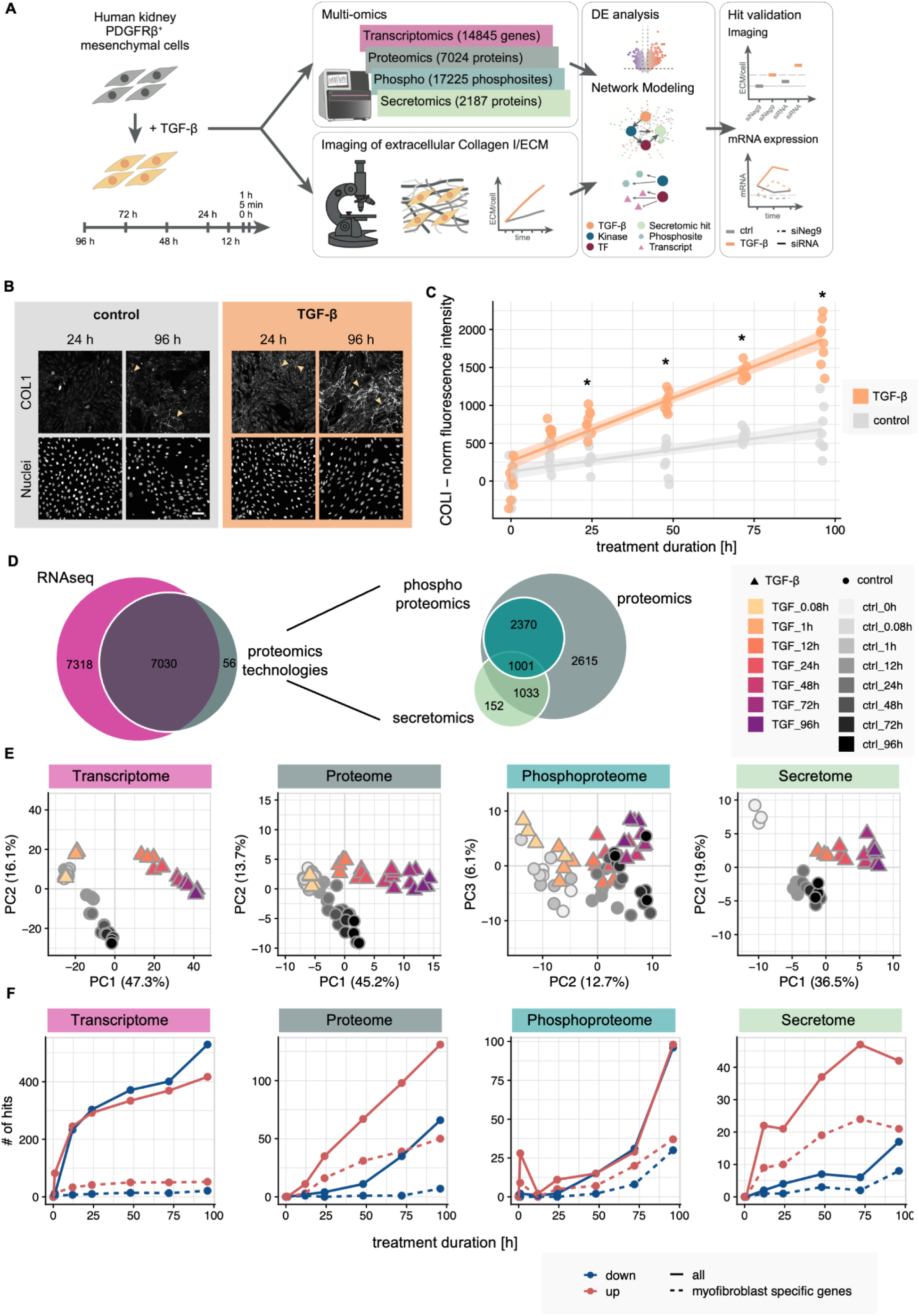
Phenotypic and molecular features defining fibrosis in PDGFRβ^+^ mesenchymal cells. (A) Overview of experimental and bioinformatic strategy. Human kidney PDGFRβ^+^ mesenchymal cells were treated with 10 ng/ml TGF-β for 5 min (0.08 h), 1, 12, 24, 48, 72, 96 h. Investigation of extracellular matrix changes over time and multi-omics data integration via mechanistic modeling. Factors of interest were further validated. (B) Immunofluorescence staining of extracellular COL1 and nuclei are shown at 96 h in control (macromolecular crowding + ascorbic acid) and TGF-β (TGF-β + macromolecular crowding + ascorbic acid) treated condition. Arrows highlight fibrillar collagen. Scale bar = 100 µm. (C) Extracellular COL1 fluorescence intensity over the specified time points and treatment conditions. Quantification of COL1 fluorescence staining was performed following background correction, correction for cell autofluorescence, and normalization to nuclei number. Single replicates are depicted by the dots (n = 36 images are averaged per replicate). Data was square root transformed for statistical analysis using a linear mixed model (* corresponds to p-value < 0.05). (D) Venn diagrams representing overlap of factors measured in the different data modalities after data processing filtering and normalization. (E) PCA scatter plots of PC1 and PC2 for the different omics modalities. For phosphoproteomics, PC2 and PC3 are depicted. The gray color scale shows control samples over time, the yellow to violet gradient resolves the time for the TGF-β-treated samples. The shape of the points shows the control samples compared to the samples treated with TGF-β. (F) Number of hits per modality and time point. Solid lines show the number of significantly deregulated transcripts (limma, absolute log2 fold-change > log2(2) and adjusted p-value < 0.05) and proteins (limma, absolute log2 fold-change > log2(1.5) and adjusted p-value < 0.05). Dashed lines show the number of significantly affected transcripts and proteins which overlap with genes specifically expressed in myofibroblasts of CKD patients (based on human PDGFRβ^+^ level 2 specificity scores by Kuppe et al. 2021).

Within 96 hours of TGF-β stimulation, we observed a robust fibrosis-like phenotype as shown by COL1 secretion (Figure 1B, Figure S1A, B), morphological changes (actin stress fiber formation, Figure S1C) and SMAD2/3 phosphorylation (Figure S2). Quantification of the extracellular COL1 fluorescence intensity revealed a significant increase of COL1 deposition upon TGF-β stimulation after 24 hours compared to the corresponding control at 24 hours, while we also observed increased collagen levels in the control conditions likely due to the macromolecular crowding agent and ascorbic acid used to accelerate collagen deposition (Figure 1C). This highlights the accelerated time scale in which a fibrosis-like phenotype can be achieved with this setup.

To gain insights into the underlying molecular mechanisms, we generated a comprehensive omics dataset spanning multiple time points after TGF-β stimulation. By combining mRNAseq with a multi-proteomics approach, more than 14,000 biomolecules could be reproducibly quantified in this system, including RNA moieties as well as intracellular, secreted and phosphorylated proteins (Figure 1D, Figure S3A). Compared to existing datasets that previously investigated TGF-β signaling or fibrosis, this multi-omics approach provides unprecedented depth and breadth of molecular insights with temporal resolution (Arif et al. 2023; D’Souza et al. 2014; Eddy et al. 2020; Lassé et al. 2023; Zhou et al. 2020a).

Stimulation of PDGFRβ^+^ mesenchymal cells with TGF-β leads to extensive perturbation of transcript and protein abundances as well as post-translational modifications (2435 molecules affected in at least one condition) that become more pronounced over time (Figure 1E, F, Figure S3B, C, Table S1). Comparing differentially expressed factors of the different time points and modalities shows good correlation at later time points, while differences at earlier time points suggest a shift in the timing of processes on transcriptome and proteome level (Figure S3E). This is reinforced by the observation that transcriptome changes in the first phases of TGF-β stimulation exceed changes at the proteome level (Figure 1F). This temporal resolution allows for a dynamic description of the fibrosis process, providing insights into the timing and progression of molecular events.

To understand the extent to which these changes are comparable to fibrotic signaling in CKD patients, and thus to assess the translational relevance of the data presented, we compared our results with scRNAseq data from healthy and CKD patient donors (Kuppe et al. 2021). A considerable proportion (48% secretomics, 36% phosphoproteomics, 32% proteome, 1% transcriptomics) of the deregulated transcripts and proteins observed in this study belong to the set of myofibroblast-specific genes identified based on the cell type specificity scores of (Kuppe et al. 2021), (Figure 1F, Figure S3D). Specifically, we observed the activation of myofibroblast-specific gene expression as the fibrotic process progresses linking long-term patient data with *in vitro* data obtained over the course of hours.

Taken together, our data provides a detailed coverage for the dynamic molecular and phenotypic changes occurring in PDGFRβ^+^ mesenchymal cells following TGF-β stimulation.

### 2.2 Functional characterization of signaling and transcription processes in kidney fibrosis

The induced molecular phenotype and its dynamic nature become particularly evident when studying specific examples over time and across different omics layers (Figure S4A, B).

Our data shows a strong upregulation of known fibrosis readouts, but with distinct temporal dynamics (Figure 2A). One example is fibronectin (FN1), a glycoprotein involved in collagen assembly and proposed drug target for fibrotic diseases (Moita et al. 2022), that shows strong upregulation over time in both transcriptome and proteome. Periostin (POSTN), a protein involved in cross-linking of the ECM, differentiation and kidney fibrosis (François & Chatziantoniou 2018; Schnieder et al. 2020), is immediately secreted upon stimulation. In contrast, Tenascin (TNC) maintains consistently high levels. TNC, an extracellular matrix glycoprotein produced by fibroblasts, promotes their activation and proliferation in an autocrine fashion (Huang et al. 2023; Moita et al. 2022). Tensin-1 (TNS1), a protein involved in integrin signaling, myofibroblast differentiation, and ECM deposition, is affected in expression as well as heavily phosphorylated which has not been reported before as a regulation mechanism in fibrosis (Figure 2A, (Bernau et al. 2017; Huang & Lo 2023). These variations underscore the differential temporal regulation of fibrotic markers in our experimental cell system.

**Figure 2:**
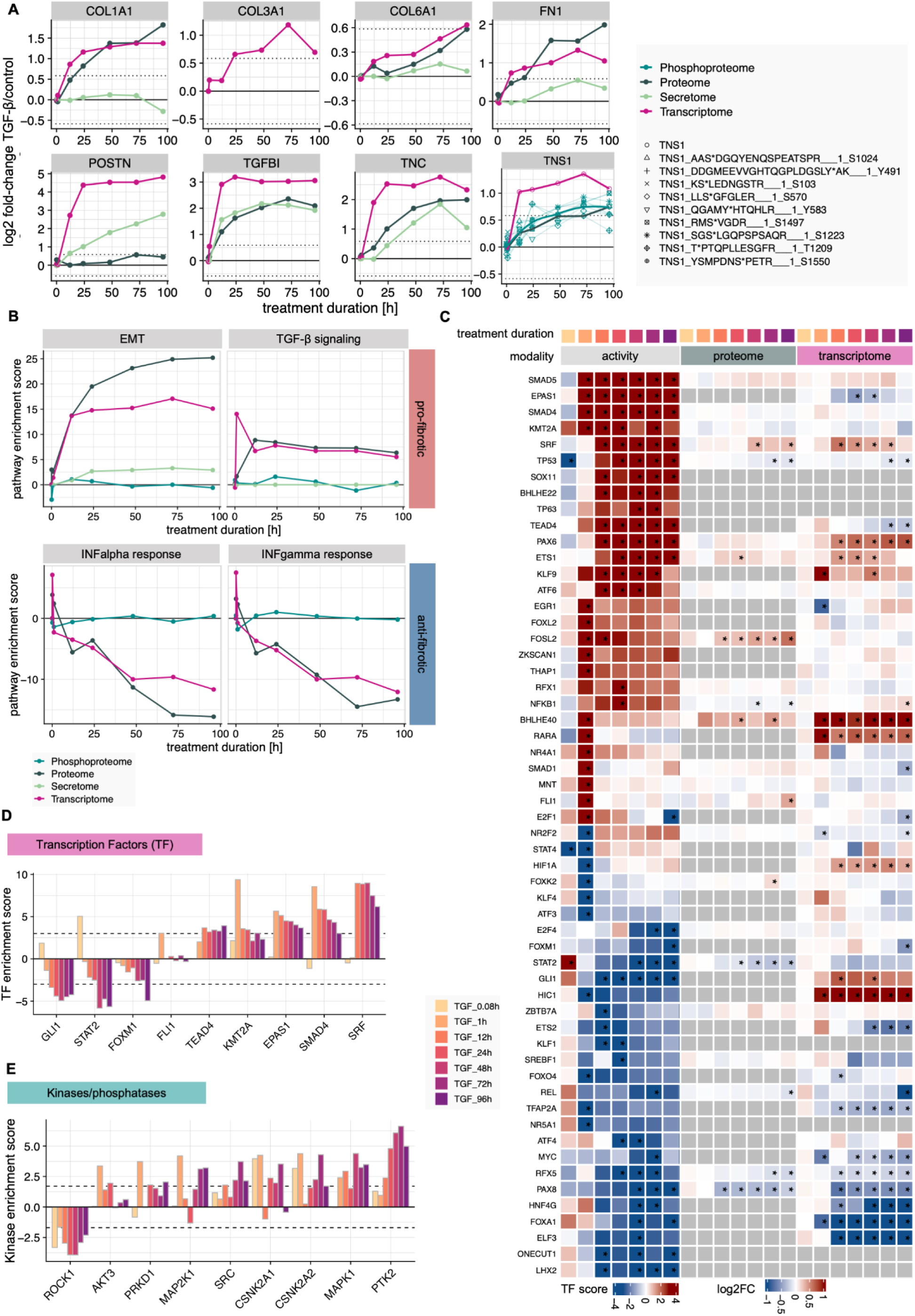
Functional characterisation of signaling and transcription processes in kidney fibrosis. (A) Differential abundance of known fibrotic markers upon TGF-β stimulation per time point in the different modalities, coded by color. (B) Selected significantly enriched pathways (GSEA using decoupleR with MSIGDB reactome database, p-value < 0.05) per time point and modality, coded by color. (C) Heatmap of differential protein and transcript abundance and matched activity scores of transcription factors per time point. Transcription factors were considered if they showed significantly altered activity upon TGF-β stimulation (enzyme activity enrichment analysis using decoupleR, p-value < 0.05 and absolute enrichment score > 3). (D) Transcription factor and (E) kinase/phosphatase enrichment scores per time point for selected significantly affected examples (enzyme activity enrichment analysis using decoupleR, p-value < 0.05 and absolute enrichment score > 3 for transcription factors or absolute enrichment score > 2 for kinases/phosphatases)

Similarly, pathways known to exert a profibrotic effect such as EMT and TGF-β are upregulated while antifibrotic pathways like interferon signaling are downregulated with distinct temporal dynamics (Figure 2B, Figure S4C, Table S2).

Illustrating these time-dependent changes, our data on COL1A1 mRNA expression supports and extends the findings from the imaging assay (Figure 1B, C, Supplementary Figure 1A, B). We observed an increase in COL1A1 mRNA and protein levels upon TGF-β stimulation over time, contrasting with a decrease in its abundance in the secretomics data. This pattern aligns with the fact that COL1A1 is incorporated into the insoluble fraction of the ECM, as suggested by previous studies (Hynes & Naba 2012; Naba et al. 2016). Furthermore, these results highlight the role of COL1 as an important regulator in the positive feedback mechanisms driving fibrosis such as influencing the matrix stiffness, demonstrating how individual components contribute to the complex, time-dependent nature of the fibrotic response observed at the pathway level (Agarwal et al. 2020; Devos et al. 2023; Kim et al. 2022; Zhou et al. 2020b).

We next inferred the activity of TFs and kinases from differential transcript and phosphoprotein abundance information, respectively, using decoupleR (Badia-I-Mompel et al. 2022). 128 transcription factors (Figure 2C, D, Table S3) and 70 kinases/phosphatases (Figure 2E, Figure S4D, Table S3) show significantly altered activity upon TGF-β stimulation with different temporal dynamics. The top hits include important drivers of TGF-beta signaling such as members of the SMAD family (Figure 2C, D). The underlying transcriptional changes of this activity prediction can be investigated by subsetting the differential expression information for known targets of a TF of interest, e.g. for SMAD4 (Figure S4E). Other interesting factors include KMT2A encoding MLL1 which has a reported role in renal fibrogenesis (Jin et al., 2022) as well as TEAD4 and SRF coactivators in YAP-TAZ signaling which leads to the production of ECM proteins in a mechanosensitive manner (Bernau et al., 2017; Hinz & Lagares, 2020; Kim et al., 2022). The consequences of JAK-STAT signaling, such as the activity of the transcription factors STAT2 and GLI1, can also be seen in this analysis, with a peak in activity after 15 minutes, followed by decreased activity but increased expression in the case of GLI1, indicating a complicated, time-dependent transcriptional regulation (Türei et al. 2016, 2021). While most of these transcription factors show a constitutive up/down regulation over time, there are examples such as FLI1 with a temporally regulated activity that only increases after one hour of stimulation (Figure 2D).

Additionally, we performed a similar analysis for kinases and phosphatases based on the obtained phosphoproteomic data (Figure 2E, Figure S4E). While the canonical response to TGF-β stimulation via SMADs is well reflected in the transcription factor activity estimation results, non-canonical fibrotic signaling pathways such as MAPK (MAPK1, MAP2K1, PTK2), RHO-ROCK (ROCK, PTK2) and PI3K-AKT (AKT3) become evident in this analysis (Figure 2E, Figure S4E) (Park & Yoo 2022). In addition, the analysis suggests that TGF-β stimulation activates less well-characterized pathways via casein kinase 2 (CSNK2A1, CSNK2A2), which could be an option for further investigation and characterization (Borgo et al. 2021).

The multi-omics analysis of TGF-β-induced kidney fibrosis revealed complex temporal dynamics in the expression and activity of fibrosis markers, signaling pathways, transcription factors, and kinases/phosphatases, demonstrating both canonical and non-canonical responses while uncovering potential novel regulatory mechanisms and therapeutic targets for further investigation.

### 2.3 Developing a time-resolved multi-omic mechanistic model of kidney fibrosis signaling

We next integrated the findings obtained from the differential expression and kinase and TF activity analyzes in a network model, using a modified version of COSMOS, an optimisation method that identifies putative causal paths explaining changes in enzymes with altered activity and multi-omics measurements based on a causal PKN (Dugourd et al. 2021).

To capture the temporal dimension of the observed fibrotic response, we created network models for both early and late stages, built from the corresponding subsets of the multi-omics data (detailed description in method section Network modeling). We focused on the obtained secretomics data to account for autocrine signaling over time by using early secreted factors as upstream input for the late network model. This provides a mechanistic molecular hypothesis for the observed ECM deposition at later time points, thus reflecting the dynamic nature of cellular communication (Figure 3A). As underlying PKN we used a directed and signed protein-protein interaction network retrieved from Omnipath (Türei et al. 2016, 2021).

**Figure 3:**
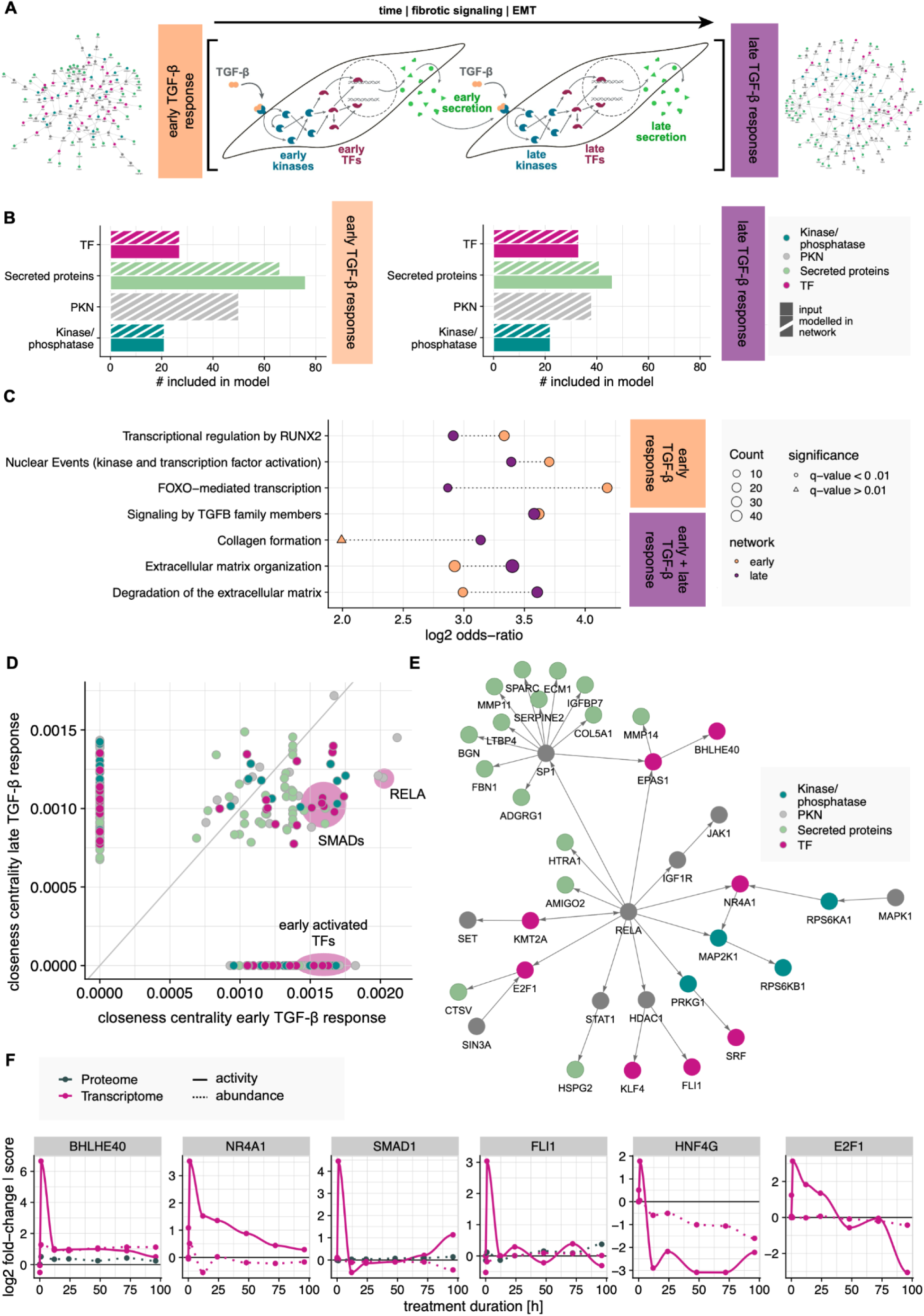
Developing a time-resolved multi-omic mechanistic model of kidney fibrosis signaling. (A) Schematic representation of the chosen computational integration strategy. Two network models were created to reflect the early and late response to TGF-β stimulation integrating transcriptomics, phosphoproteomics and secretomics data for different time points. By considering early secreted factors as upstream stimuli for the late response network, autocrine signaling can be modeled. To generate the network models the COSMOS method was used with a PKN consisting of signed directed protein-protein interactions retrieved from Omnipath. (B) Properties of the resulting network models. For both models, all input enzymes (TFs and kinases/phosphatases) as well as the majority of considered secreted proteins could be integrated in the solution. (C) Selected significantly overrepresented pathways in the early and late network models (pathway overrepresentation analysis of network nodes using the Reactome pathway database, p-value < 0.05). Point color indicates the network for which the enrichment has been calculated. Point size shows the number of nodes in the network mapped to the pathway and point shape the significance (q-value). (D) Comparison of closeness centrality measures per node for the early and late TGF-β response network model. Points are colored by node type. (E) RELA related signaling in the early TGF-β response network model. Node color indicates node type. (F) Temporal activity and abundance profiles for selected early activated TFs. The color indicates the underlying modality, the line type and the data type.

Besides proteins with altered secretion upon TGF-β stimulation, we selected the observed early and late deregulated TFs and kinases/phosphatases as input nodes and confirmed that all moieties are represented in the PKN (Figure S5A). The integration process incorporated all input enzymes and measurements, as well as most secreted proteins used as model input, while maintaining a manageable solution size (Figure 3B, Figure S5B, Table S4, Table S5).

The resulting networks reflect the expected TGF-β response, as shown by an overrepresentation analysis of the network nodes (Figure 3C, Figure S5C). TGF-β signaling and ECM remodeling processes are among the most enriched pathways in both models, while a series of effects on transcriptional regulation seem to be strengthened in the early network. The signature of TGF-β signaling in the networks is extensive, linking several TFs, kinases and secreted proteins highlighting the value of integrating different modalities (Figure S5D).

Analyzing node connectivity based on their closeness centrality in the network revealed that transcription factors and kinases are more connected than secreted proteins (Figure 3D, Figure S5E), with for example SMAD TFs remaining critical in both early and late networks. While this pattern may derive from prior knowledge, a number of network-exclusive nodes as well as nodes with different closeness centrality between the two networks illustrate shifts in the connectivity of the nodes and thus potentially biological relevance over time (Figure 3D).

As mentioned before, network node enrichment analyses underscored the significance of early transcriptional events, while the analysis of node centrality underscores the importance of the TF RELA in the early network (Figure 3C, D, E). RELA is being studied in the context of fibrotic disease (Chung et al. 2017; Moles et al. 2013) and is connected to many early-activated TFs in the network, some well-documented in fibrosis (Goffin et al. 2010; Kendall & Feghali-Bostwick 2014; Kum et al. 2007; Mikhailova et al. 2023).

A group of TFs governed by RELA showed a similar activity pattern characterized by an early activation peak (Figure 3E, F). We focused on a number of these early-acting TFs that have been poorly or not at all characterized in the context of fibrosis, but were predicted to have a regulating role on differential mRNA expression. Those include Transcription Factor E2F1, Friend leukaemia integration 1 transcription factor (FLI1), Nuclear Receptor Subfamily 4 Group A Member 1 (NR4A1), and Class E basic helix-loop-helix protein 40 (BHLHE40) for validation, which are all predicted to be downstream of RELA. Similar profiles, with an upregulated activity at 1 h post TGF-β stimulation were shown by the TFs Mothers against decapentaplegic homolog 1 (SMAD1), and Hepatocyte nuclear factor 4-gamma (HNF4G).

To summarize, the integration of multi-omic data into time-resolved network models of early and late fibrotic responses revealed dynamic shifts in signaling pathways, transcription factor activities, and protein interactions, highlighting the temporal complexity of kidney fibrosis progression and identifying both well-known and novel regulatory factors for further investigation.

### 2.4 Experimental validation confirms the implication of selected TFs in regulating ECM deposition

To further validate the role of these transcription factors in the development of fibrotic diseases, we exploited the perturbability of the used *in vitro* model system. We investigated their role in ECM remodeling and collagen deposition using siRNA knockdowns and subsequent imaging of the deposited ECM. By monitoring ECM production, we took advantage of a direct phenotypic readout of the knockdown performed, while RT-qPCR experiments allowed us to track the effects of TF knockdowns on the mRNA expression of their targets at the molecular level (Figure 4A, Table S9).

**Figure 4:**
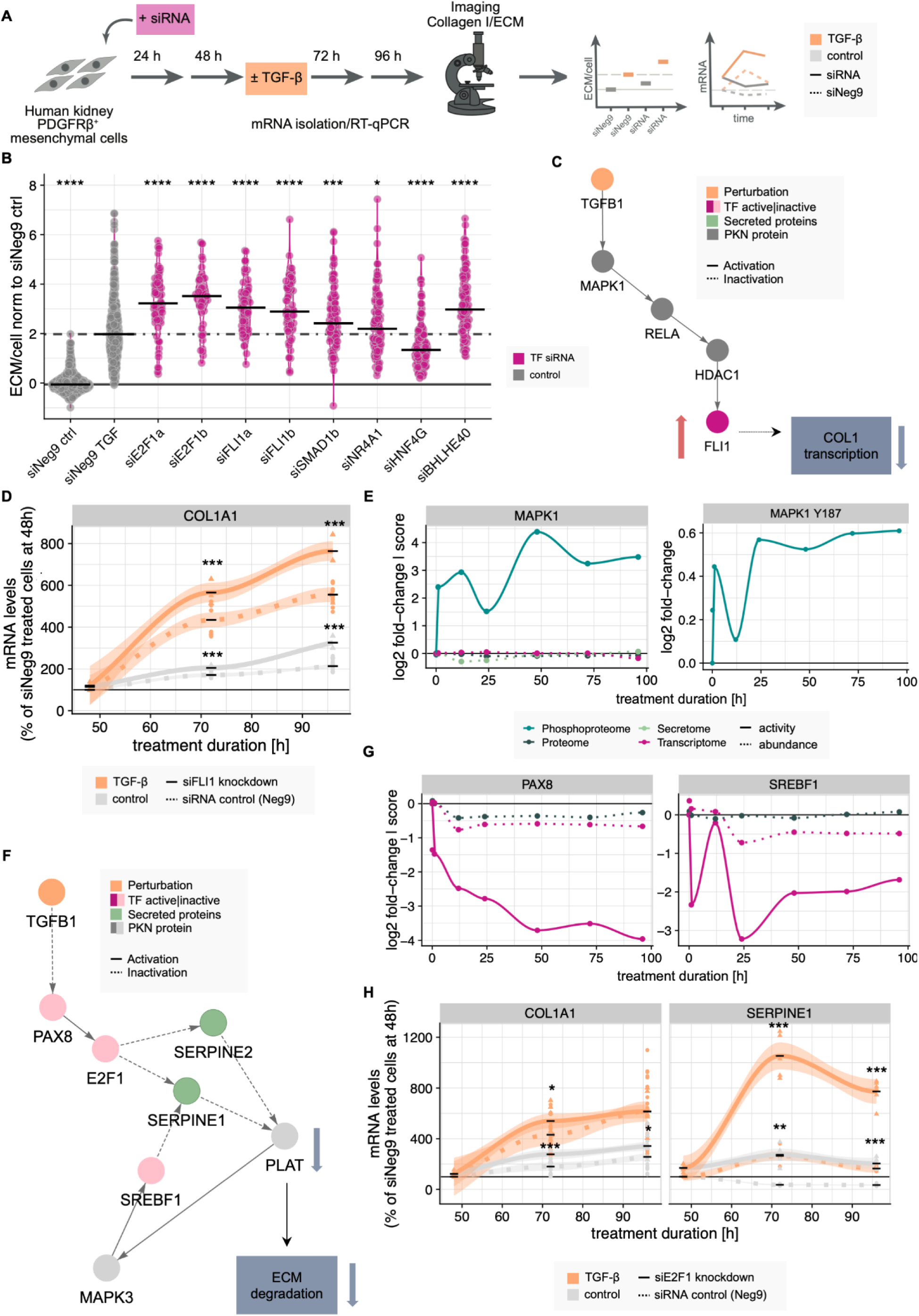
Validation of predicted fibrosis modulators and related molecular mechanisms. (A) Schematic representation of the experimental workflow for validation experiments. (B) Fluorescence intensity of ECM for selected TF knockdowns (purple, 96h knockdown followed by 48h TGF-β treatment) and the siNeg9 TGF-β treated/unstimulated controls (gray). Intensities of each knockdown were compared to the siNeg9 TGF-β treated condition (siNEg9 TGF) using a unpaired two-sided t-test (* corresponds to p-value < 0.05, ** corresponds to p-value < 0.01, *** corresponds to p-value < 0.001). Black bars represent the median fluorescence intensity per distribution. (C) Subnetwork of early TGF-β response network model showing directed protein-protein interactions leading to FLI1 activation. Node color indicates node type, edge line type indicates activation/inhibition. (D) RT-qPCR data to confirm FLI1 knockdown effect on its potential downstream target COL1A1 +/-TGF-β stimulation at different time points. Color indicates TGF-β stimulation vs control, line type indicates siNeg9 control vs TF knockdown. Significance has been tested per time point and treatment condition using a t-test (* corresponds to p-value < 0.05, ** corresponds to p-value < 0.01, *** corresponds to p-value < 0.001). (E) Temporal activity and abundance profiles for the MAPK1 kinase and the MAPK1 Y187 phosphorylation site. The color indicates the underlying modality, the line type shows the data type. (F) Subnetwork of late TGF-β response network model showing directed protein-protein interactions leading to E2F1 inactivation upstream of SERPINE1 activation. Node color indicates node type, edge line type indicates activation/inhibition. (G) Temporal activity and abundance profiles for the PAX8 and SREBF1 transcription factors. The color indicates the underlying modality, the line type shows the data type. (H) RT-qPCR data to confirm the effect of E2F1 knockdown on its potential downstream targets COL1A1 and SERPINE1 +/-TGF-β stimulation at different time points. Color indicates TGF-β stimulation vs control, line type indicates siNeg9 control vs TF knockdown. Significance has been tested per time point and treatment condition using a two-sided unpaired t-test (* corresponds to p-value < 0.05, ** corresponds to p-value < 0.01, *** corresponds to p-value < 0.001).

Analysis of the fluorescence intensity of the deposited ECM after knockdown of the early activated TFs revealed a general trend towards increased deposition compared to a siNeg9 control (Figure 4B, Figure S6A). This effect is considerably pronounced upon TGF-β stimulation, indicating the importance of these TFs as potential negative regulators of collagen secretion given fibrotic signaling (Figure 4B, Figure S6A right panel). All six TF knockdowns had a significant impact on ECM deposition (unpaired two-sided t-test, p-value < 0.05), but for NR4A1 the effect magnitude was quite small, while the HNF4G knockdown was the only example that showed an opposite effect. Notably, the activity of HNF4G was strongly downregulated after 12 hours of TGF-β stimulation, which is not the case for the other TFs we considered for validation (Figure 3F). The collagen knockdowns, performed as a positive control, led to the expected reduction in collagen deposition (Figure S6A). Interestingly, the varying effects of SMAD1 knockdown on ECM deposition levels (Figure S6A) could be attributed to differences in knockdown efficiency between the two siRNAs used (Figure S6B).

By combining insights into potential mechanisms of TF activity from our multi-omics models with information on the expression of target genes upon knockdown, we can investigate the role of specific TFs in detail. FLI1 is involved in early signaling around RELA, with an activity peak at 1 h and no deregulation of activity or abundance thereafter (Figure 3E, F). Further investigation of the network reveals a possible regulation via MAPK1-RELA-HDAC, which is downstream of TGFB1 and leads to the activation of FLI1 (Figure 4C). Consistent with the imaging data, COL1A1 mRNA production is also increased upon FLI1 knockdown on mRNA level (Figure 4D, Figure S6C, D, Table S10). These observations align with existing literature describing the role of FLI1 as an inhibitor of collagen production controlled by HDAC1 (Figure 4C) (Mikhailova et al. 2023; Wang et al. 2006). Regulation by MAPK1 is further supported by the predicted increased activity of the kinase at an early time point, as well as significantly increased phosphorylation in the activation loop of the kinase (Y187, limma, adjusted p-value < 0.05, absolute log2 fold-change > log2(1.5)), which has been reported to lead to its increased activity (Figure 4E, (Schmidlin et al. 2019). These findings highlight the benefits of the integrative approach, as they would not have been detected by a single omics modality.

A second very interesting case is the transcription factor E2F1. In this case, activity estimation predicts activity at earlier time points, and increased ECM deposition is observed in images after knockdown, as in most other early-activated TFs (Figure 3F, Figure 4B). Interestingly, E2F1 activity is not significantly affected between 12 and 72 hours, but at the last monitored time point (96 hours), the expression of the E2F1 targets and thus its activity decreases substantially (Figure 3F). E2F1 is included downstream of RELA in the early network (Figure 3E) and in the late network model, in which it is predicted to be downregulated in its activity downstream of inactive PAX8, which in turn could be inactivated by TGFB1 signaling (Figure 4F). These observations are consistent with the estimates of TF activity for PAX8, which shows a strong down-regulation of TF activity over time (Figure 4G). This suggests that the activity shift in early activated TFs is driven by complex transcriptional reprogramming in response to TGF-β stimulation. Moreover, E2F1 regulation is associated with secretory processes, as shown by the network model, which suggests that SERPINE1 (also known as plasminogen activator inhibitor-1 PAI-1) and SERPINE2 secretion, key modulators of collagen deposition, are upregulated downstream of E2F1 inhibition (Figure 4F, Figure S6E). SERPINE1 and SERPINE2 do not cause this effect by influencing COL1 expression, but by inhibiting collagen degradation and thus contributing to increased extracellular matrix deposition (Bergheim et al. 2006; Qi et al. 2008). This effect is confirmed at the SERPINE1 mRNA level, which increases considerably upon E2F1 knockdown (Figure 4H, Figure S6F, G). Furthermore, the RT-qPCR data supports the hypothesis that E2F1 has an indirect effect on collagen deposition, as the expression of COL1A1 mRNA is only moderately affected by E2F1 knockdown, in contrast to the observations for FLI1 (Figure 4H, Figure S6F, G).

In summary, the integration of extensive time-resolved multi-omics data into mechanistic networks in combination with phenotypic and molecular validation experiments can serve as a basis to develop working models for molecular mechanisms underlying kidney fibrosis, as demonstrated in this study for FLI1 and E2F1.

## 3. Discussion

In this study, we present an integrative approach to investigate the complex molecular mechanisms underlying kidney fibrosis. By combining a perturbable human PDGFRβ^+^ mesenchymal cell *in vitro* model system with time-resolved multi-omics profiling and advanced computational analyses, we have generated a comprehensive dataset that provides unprecedented insights into the dynamic nature of fibrotic processes driven by these cells.

Our *in vitro* model system enables detailed phenotypic and molecular characterization of fibrosis in PDGFRβ^+^ mesenchymal cells, addressing many limitations of existing approaches. The ability to observe and quantify phenotypic consequences of TGF-β stimulation, such as COL1/ECM deposition within a 96-hour timeframe demonstrates the accelerated nature of our system. This rapid induction of a fibrotic phenotype allows for more efficient studies of potential therapeutic interventions and enables us to link long-term patient data, spanning years, with *in vitro* data obtained over the course of hours. This association between *in vitro* and *in vivo* data strengthens the potential of our model system for identifying clinically relevant therapeutic targets.

The multi-omics approach, encompassing transcriptomics, proteomics, phosphoproteomics, and secretomics, has allowed us to quantify over 14,000 biomolecules across multiple time points, of which 2,435 were significantly affected in at least one condition. The temporal resolution of our data has uncovered distinct dynamics in the expression and activity of known and potentially novel biomarkers and modulators of fibrosis, highlighting the importance of time-dependent analyses in understanding disease mechanisms.

A key strength of our approach is the ability to distinguish between abundance and activity of molecular players. This distinction is particularly evident in our analysis of transcription factor and kinase activities, which has revealed novel insights into the regulatory networks driving fibrosis.

Our multi-omics approach, combined with network modeling and experimental validation, revealed critical regulatory mechanisms that single-omics studies might overlook. Perturbation experiments of our *in vitro* model system through siRNA knockdown experiments allowed us to validate the computational predictions and explore the functional roles of specific factors in fibrosis. An example of this is the unexpected finding that knockdown of several early-activated transcription factors such as E2F1, FLI1, SMAD1, NR4A1, HNF4G, and BHLHE40 leads to increased collagen deposition. This further suggests that these factors may act as negative regulators of fibrosis, thereby opening new avenues for therapeutic intervention.

While our study provides valuable insights, it also has limitations. Despite the power of our integrative approach, there are still aspects that we do not fully understand, such as the precise mechanisms causing the downregulation of certain transcription factors. Additionally, our network model provides valuable insights and potential mechanisms, but these need to be thoroughly validated.

Furthermore, while our temporal profiling provides a detailed view of early fibrotic events, longer-term studies may be necessary to fully understand the chronic nature of kidney fibrosis. To move forward, the impact of our results should be confirmed with more patient data, bridging the gap between our *in vitro* findings and clinical observations.

The *in vitro* nature of our model system may not fully recapitulate the complex multicellular interactions present in the kidney. Future studies could address this by incorporating co-culture systems or organoid models to better reflect the *in vivo* environment.

In conclusion, our integrative, time-resolved multi-omics approach provides a comprehensive view of the molecular events driving kidney fibrosis. By combining advanced experimental and computational methods, we have generated a rich resource for the renal research community and demonstrated the power of systems biology approaches in unraveling complex disease mechanisms. The insights gained from this study not only advance our understanding of kidney fibrosis but also pave the way for the development of novel therapeutic strategies targeting this challenging condition. Future work leveraging the comprehensive data of this study has the potential to impact the clinical management of chronic kidney disease and other fibrotic disorders.

## 4. Materials and Methods

LLM (Claude 3.5 Sonnet and ChatGPT-4) were used for formulations of text and editing.

### 4.1 Cell Biology

#### Cell Lines and Reagents

Human kidney PDGFRβ^+^ mesenchymal cells were received from the Kramann lab (for further information please check (Kuppe et al. 2021)) and cultured in low glucose DMEM growth medium (Gibco 31885) supplemented with 5% FBS (Gibco A5256701). Cells were maintained at 37°C in a humidified incubator with 5% CO2 and passaged approximately three times a week. Mycoplasma testing was routinely conducted, yielding negative results.

#### Experimental Setup

For multi-omics and time point experiments, the medium in all conditions was changed 24 hours post-plating. The longest time point treatment (e.g. 96 hours) was initiated the day after seeding. Control samples were maintained in low glucose DMEM without phenol red (Gibco 11880), supplemented with 1% L-Glutamine (Sigma Life Science G7513), Ficoll 70 and 400 (Sigma Aldrich F2878 and F4375), and 500 μM L-ascorbic acid 2-phosphate (Cayman Chemical Company 16457). TGF-β treated samples received the same medium with an additional 10 ng/ml of recombinant human TGF-β1 (R&D systems 240-B-010). Until the treatment started and for the 0 h treatments, the cells were kept in DMEM without phenol red, FBS, Ficoll or ascorbic acid. Media changes were performed daily, with specific treatments applied as described to ensure all samples were ready for downstream processing at the same time.

#### siRNA Transfections

All siRNAs used are described in Table S7 and were acquired from Ambion/Thermo Fisher. Cells were plated one day before siRNA transfections, targeting 40-50% confluency on the day of treatment. The ScreenFect®siRNA protocol was employed. For a single 24-well plate reaction, 1 μl of ScreenFect®siRNA reagent (ScreenFect S-4001) was mixed with 39 μl of Dilution Buffer (ScreenFect S-2001). Separately, 5 pmol siRNA was diluted with 39 μl of Dilution Buffer. The mixtures were combined, incubated for 20 minutes at room temperature, and then 420 μl of fresh DMEM (without FBS) was added. The transfection mixture replaced the cell medium, which was changed to DMEM + 5% FBS after 5-6 hours. For knockdown and TGF-β treatment experiments, cells were cultured for 48 hours post-siRNA transfection in DMEM + 5% FBS before starting TGF-β treatments, with media changes every 24 hours. Cells were fixed/harvested at specified time points

#### Immunofluorescence Assays

Cells were seeded on glass-bottom 24-well plates (Cellvis P24-1.5H-N) and treated as described above. After aspirating the medium, cells were washed with PBS and fixed with 4% PFA (Thermo Fisher Scientific 50-980-491) containing 1:1000 Hoechst 33342 (Thermo Fisher Scientific H21492) for 10-15 minutes at room temperature. Following three PBS washes, cells were used for immunofluorescence staining.

#### Extracellular Immunofluorescence Staining

For extracellular matrix (ECM) visualization, cells were incubated with anti-COL1 antibody (Rockland 600-401-103-0.5, 1:500 in PBS) for 1-1.5 hours at room temperature, washed, and then incubated with fluorescently labeled secondary anti-rabbit IgG AlexaFluor 488 (Molecular Probes A11008, 1:400 in PBS) in PBS for 30-45 minutes. Washed cells were kept in PBS and imaged. Due to issues with new batches of the anti-COL1 antibody, GFP-labeled CNA35 dye (EMBL protein expression facility, 1:250 in PBS) was used for validation experiments (siRNA knockdowns of TFs). After fixation and washing, cells were incubated with CNA35 for 1-1.5 hours, washed, and imaged. In cases of increased autofluorescence from siRNA transfection, cells were stained with an anti-GFP (Origene TP401) followed by Alexa 647-conjugated secondary anti-rabbit (Invitrogen A21245).

#### Intracellular Immunofluorescence Staining

Intracellular proteins, such as actin, were visualized by permeabilizing cells with 0.1% Triton X-100 (Sigma Aldrich T8787) for 15 minutes post-fixation. Subsequent incubation with Phalloidin AlexaFluor 647 (Invitrogen A22287), washing, and confocal imaging were performed as described for extracellular staining.

### 4.1 Microscopy

#### Wide-field Microscopy

High-throughput imaging was conducted using the Molecular Devices IXM automated widefield screening microscope. A total of 36 fields of view were captured per well using a CFI P-Apo 20x Lambda/0.75 objective. Nuclear signals were acquired in the Hoechst channel (Ex: 377/50, Em: 477/60), with ECM signals in the GFP channel (Ex: 472/30, Em: 520/35) or other channels depending on the secondary antibody used (such as Cy5 with Ex:28/40-25 Em: 692/40-25).

#### Confocal Microscopy

Confocal microscopy was performed using a Zeiss LSM 900 microscope. For visualization of cell cytoskeleton changes (actin filaments), Z-stacks were acquired covering the entire cell thickness, with parameters and number of Z-stacks kept constant across samples. The microscope was equipped with a Plan-Apochromat 20x/0.8 M27 air objective (FWD=0.55mm). Three lasers were utilized: 405 nm (5 mW), 488 nm (10 mW), and 640 nm (5 mW). For detection, a Gallium Arsenide Phosphide-PMT (GaAsP-PMT) was used for fluorescence. The filter configuration included excitation filters BP 385/30 for DAPI/Hoechst, BP 469/38 for FITC/Alexa Fluor 488, and BP 631/33 for Cy5. A QBS 405+493+575+653 beam splitter was employed, along with an emission filter QBP 425/30+514/30+592/25+709/100.

### 4.1 Biochemistry

#### Cell Lysis and Sample Preparation

Cells were washed twice with ice-cold PBS and lysed in Pierce RIPA buffer (Thermo Fisher Scientific 89900) with EDTA-Free Protease Inhibitor Cocktail (Roche 1836170001) for 5 min on ice. Lysates were centrifuged at 14,000 g for 15 min at 4°C. Supernatants were used for analysis or stored at −80°C. Protein samples were mixed with 2x sample buffer (200 mM Tris-HCl (Sigma-Aldrich T5941), 25% glycerol (v/v) (Sigma-Aldrich 15523-1L-R), 11.25% SDS (v/v) (Bio-Rad Laboratories 1610394), 325 mM DTT (Sigma-Aldrich D0632), 0.0125% (w/v) bromophenol blue (Sigma-Aldrich B5525), pH 6.8) and heated at 98°C for 5 min.

#### SDS-PAGE and Western Blot

Protein samples were separated on NuPAGE 4-12% Bis-Tris gels (Thermo Fisher Scientific NP0321, NP0322) using NuPAGE MOPS SDS running buffer (Thermo Fisher Scientific NP0001) at 90-100 V for 200 min. Proteins were transferred to Immobilon PVDF membranes (pore size 0.45 μM, Merck Millipore IPVH00010) for 1 h at 100 V. Membranes were blocked with 5% BSA (Sigma Aldrich A2153) in TBS-T (ThermoScientific J60764.K2)), were incubated with primary antibodies (Tubulin-α, Thermo Scientific MS-581-P0; SMAD2, Proteintech 12570-1-AP; phospho-SMAD2, Cell Signaling 18338) 1 h at room temperature or overnight at 4°C. After washing, the membranes were treated with HRP-coupled secondary antibodies (Anti-mouse IgG, Sigma-Aldrich A9044; Anti-rabbit IgG, Sigma-Aldrich A0545) for 45 min at room temperature. Proteins were visualized using Pierce^TM^ ECL Plus Western Blotting Substrate (Thermo Scientific 32132) and imaged with Azure 280 (Biozyme) or Bio-Rad imager. Bands were quantified using Fiji ImageJ and normalized to loading controls.

#### RNA Isolation and RT-qPCR

RNA was extracted using the RNeasy Mini kit (Qiagen 74104) according to manufacturer’s instructions. RNA concentration and purity were measured with Nanodrop 8000 Spectrophotometer (Thermo Fisher Scientific). For RT-qPCR, 300-500 ng total RNA was reverse transcribed using SuperScript IV Reverse Transcriptase (Thermo Fisher Scientific 18090200). cDNA was diluted 1:10 and amplified using SYBR Green PCR Master Mix (Applied Biosystems by Thermo Fisher Scientific 4309155) on StepOne Real-Time PCR system (Thermo Fisher Scientific) or QuantStudio 6 Flex systems (Bio-Rad Laboratories).

Data was analyzed using the 2^-ΔΔCT^ method (Livak & Schmittgen 2001), normalizing to GAPDH. Primer sequences are described in Table S6.

### 4.2 Multi-omics Experiment

#### RNAseq

Cell lysis and RNA isolation were performed using the Total RNA Purification Kit (Norgen Biotek Corporation 17200) according to the manufacturer’s protocol. On-column DNA removal was conducted using Norgen’s RNase-Free DNase I Kit (Norgen Biotek Corporation 25710) following the manufacturer’s instructions.

The initial RNA was QCed using Agilent Bioanalyzer with the RNA Nano Assay kit as per the manufacturer’s protocol. The RNA sample set was then standardized to 300 ng total RNA in 50 µl using the concentration values given by the Bioanalyzer. The libraries were prepared on a Beckman Coulter Automated Workstation Biomek i7 Hybrid (MC +Span-8). For library preparation an automated version of the NEBNext® UltraTM II Directional RNA Library Prep Kit was used, following section 1 - Protocol for use with NEBNext Poly(A) mRNA Magnetic Isolation Module. An adaptor dilution of 1 to 20 was used, the samples were individually barcoded using unique dual indices during the PCR using 13 PCR cycles as per the manufacturer’s protocol. The individual libraries were quantified using the Qubit HS DNA assay as per the manufacturer’s protocol. For the measurement 1ul of sample in 199ul of Qubit working solution was used. The quality and molarity of the libraries was assessed using Agilent Bioanalyzer with the DNA HS Assay kit as per the manufacturer’s protocol. The assessed molarity was used to equimolarly combine the individual libraries into one pool for sequencing. The pool was loaded and sequenced on an Illumina NextSeq 2000 platform (Illumina, San Diego, CA,USA) using a P3 50 cycle kit, a read-length of 72 bp single-end reads and 650 pM final loading concentration.

#### Proteomics experiments

Proteomics samples were lysed on-plate and processed by the Proteomics Core Facility.

##### Sample preparation

For all samples, protein concentration was determined using the Pierce BCA Protein Assay Kit (Thermo Fisher Scientific 23227). Working reagent and BSA standards were prepared followed by the manufacturer’s protocol. 25 μl of standards and 2 μl of TCA-precipitated samples were added to NuclonTM Delta Surface 96-well plate (Thermo Fisher Scientific 167008) in triplicates. 200 μl of working reagent was added to each well. Plates were incubated at 37°C for 30 min after 1 min shaking. Absorbance was measured at 562 nm using a Infinite M1000 pro plate reader (Tecan). Protein concentrations were calculated using a standard curve, and samples were diluted to achieve equal protein amounts across all samples.

For the secretomics experiments, cell culture supernatants were collected and centrifuged at 300 g for 5 min at 4°C to remove cells. Cleared supernatants were snap-frozen and stored at −80°C until further processing. Proteins were enriched using TCA precipitation. Therefore, one part ice-cold TCA (Merck 1008071000) was added to four parts of the protein sample. Samples were subsequently vortexed and incubated on ice for 20-30 min. This was followed by centrifugation at 10,000 g for 20 min at 4°C. Pellets were washed with 500 μl ice-cold 10% TCA, vortexed, and centrifuged at max speed for 20 min at 4°C. A final wash was performed with 1 ml ice-cold acetone (−20°C) (Merck 100014), followed by centrifugation at max speed for 30 min at 4°C. Dried pellets were dissolved in 25 μl 1% SDS buffer (1% SDS (Bio-Rad 1610418), 50 mM HEPES (Gibco 15630-080) in DEPC-treated water (Ambion AM9906)). Reduction of disulphide bridges in cysteine containing proteins was performed with dithiothreitol (56°C, 30 min, 10 mM in 50 mM HEPES, pH 8.5). Reduced cysteines were alkylated with 2-chloroacetamide (room temperature, in the dark, 30 min, 20 mM in 50 mM HEPES, pH 8.5). Samples were prepared using the SP3 protocol (Hughes et al. 2014, 2019) and trypsin (sequencing grade, Promega) was added in an enzyme to protein ratio 1:50 for overnight digestion at 37°C. Next day, peptide recovery in HEPES buffer by collecting supernatant on magnet and combining with second elution wash of beads with HEPES buffer. Peptides were labeled with TMT11plex (Werner et al. 2014)

Isobaric Label Reagent (ThermoFisher) according to the manufacturer’s instructions. Samples were combined for the TMT11plex and for further sample clean up an OASIS® HLB µElution Plate (Waters) was used. Offline high pH reverse phase fractionation was carried out on an Agilent 1200 Infinity high-performance liquid chromatography system, equipped with a Gemini C18 column (3 μm, 110 Å, 100 x 1.0 mm, Phenomenex) (Reichel et al. 2016).

For the full proteome and phosphoproteome experiments, phosphopeptide enrichment was essentially done as described in Potel et al. 2018 (Potel et al. 2018). Cells were lysed on the plate with 500 µl lysis buffer composed of 100 mM Tris-HCl pH 8.5, 7 M Urea, 1% Triton, 5 mM Tris(2-carboxyethyl)phosphin-hydrochlorid, 30 mM chloroacetamide, 10 U/ml DNase I (Sigma-Aldrich), 1 mM magnesium chloride, 1 mM sodium orthovanadate, phosphoSTOP phosphatase inhibitors (Sigma-Aldrich) and complete mini EDTA-free protease inhibitors (Roche). The samples were sonicated 3 times for 10 sec (continuous pulse, 50% duty cycle) with intervals of cooling on ice for 60s until the viscosity was reduced with an ultrasonic sonifier (Branson). Residual cell debris was removed by centrifugation at 17000g at 8 C for 10 min. To the supernatant 1% benzonase (Merck Millipore) was added and incubated at RT for 1 h. Then methanol/chloroform precipitation was performed by adding to 1 volume of sample 4 volumes of methanol, 1 volume of chloroform and 3 volumes of ultrapure water. Centrifugation was performed for 15 min at 4000g. The upper layer was removed without disturbing the interface and 3 volumes of methanol were added before another centrifugation step. The liquid phase was removed and the white protein precipitate was allowed to air dry. Proteins were resuspended in digestion buffer composed of 100 mM Tris-HCl pH 8.5, 1% sodium deoxycholate (Sigma-Aldrich), 5 mM Tris(2-carboxyethyl)phosphin-hydrochlorid and 30 mM chloroacetamide. Trypsin was added to a 1:50 ratio (w/w) and protein digestion was performed overnight at RT. The next day, digestion was stopped by the addition of TFA to a final concentration of 1% in the sample. The sodium deoxycholate was precipitated for 15 min at RT and the samples were centrifuged for 10 min at 17000g at RT. The supernatant was desalted by using Oasis HLB 96-well plates 30 µM (Waters). Thereby buffer A was composed of MS-grade water (Chemsolute) with 0.1% formic acid and buffer B 80% acetonitrile (Chemsolute) in MS-grade water with 0.1% formic acid. Eluted peptides were dried in a vacuum centrifuge.

For phosphopeptide enrichment, peptides were taken up in IMAC loading solvent (70% acetonitrile, 0.07% TFA). Of each sample a small aliquot was used for full proteome analysis. Phosphopeptide enrichment was performed on an UltiMate 3000 RSLC LC system (Dionex) using a ProPac IMAC-10 Column 4 x 50 mm, P/N 063276 (Thermo Fisher Scientific). To enable post-enrichment TMT labeling, the phosphopeptides were eluted with 0.4% dimethylamine (Sigma-Aldrich). Peptides were labeled with TMT16plex (Thompson et al. 2019) Isobaric Label Reagent (ThermoFisher) according to the manufacturer’s instructions. In short, 0.8mg reagent was dissolved in 42ul acetonitrile (100%) and 4 µl of stock was added and incubated for 1 h room temperature. Followed by quenching the reaction with 5% hydroxylamine for 15 min. RT. Samples were combined and for further sample clean up an OASIS® HLB µElution Plate (Waters) was used. The TMT-labelled phosphoproteome and full proteome were fractionated by high-pH reversed-phase carried out on an Agilent 1200 Infinity high-performance liquid chromatography system, equipped with a Gemini C18 column (3 μm, 110 Å, 100 x 1.0 mm, Phenomenex). 48 fractions were collected along with the LC separation that were subsequently pooled into 12 fractions. Pooled fractions were dried under vacuum centrifugation and reconstituted in 10 μL 1% formic acid, 4% acetonitrile and then stored at −80 °C until LC-MS analysis.

##### Mass spectrometry measurements

For the secretomics experiments, an UltiMate 3000 RSLC nano LC system (Dionex) fitted with a trapping cartridge (µ-Precolumn C18 PepMap 100, 5µm, 300 µm i.d. x 5 mm, 100 Å) and an analytical column (nanoEase™ M/Z HSS T3 column 75 µm x 250 mm C18, 1.8 µm, 100 Å, Waters). Trapping was carried out with a constant flow of 0.05% trifluoroacetic acid at 30 µL/min onto the trapping column for 6 minutes. Subsequently, peptides were eluted via the analytical column with a constant flow of solvent A (0.1% formic acid, 3% DMSO in water) at 0.3 µL/min with increasing percentage of solvent B (0.1% formic acid, 3% DMSO in acetonitrile). The outlet of the analytical column was coupled directly to a Fusion Lumos (Thermo) mass spectrometer using the Nanospray Flex™ ion source in positive ion mode. The peptides were introduced into the Fusion Lumos via a Pico-Tip Emitter 360 µm OD x 20 µm ID; 10 µm tip (CoAnn Technologies) and an applied spray voltage of 2.4 kV. The capillary temperature was set at 275°C. Full mass scan was acquired with a mass range 375-1500 m/z in profile mode in the orbitrap with resolution of 120000. The filling time was set at a maximum of 50 ms with a limitation of 4×105 ions. Data dependent acquisition (DDA) was performed using quadrupole isolation at 0.7 m/z, the resolution of the Orbitrap set to 30000 with a fill time of 94 ms and a limitation of 1×105 ions. A normalized collision energy of 36 was applied. MS2 data was acquired in profile mode.

For the phosphoproteomics experiments, an UltiMate 3000 RSLC nano LC system (Dionex) fitted with a trapping cartridge (µ-Precolumn C18 PepMap 100, 5µm, 300 µm i.d. x 5 mm, 100 Å) and an analytical column (nanoEase™ M/Z HSS T3 column 75 µm x 250 mm C18, 1.8 µm, 100 Å, Waters) was coupled to a Orbitrap Fusion™ Lumos™ Tribrid™ Mass Spectrometer (Thermo). Peptides were concentrated on the trapping column with a constant flow of 0.05% trifluoroacetic acid at 30 µL/min for 6 minutes. Subsequently, peptides were eluted via the analytical column using a binary solution system at a constant flow rate of 0.3 µL/min. Solvent A consists of 0.1% formic acid in water with 3% DMSO and solvent B of 0.1% formic acid in acetonitrile with 3% DMSO. The percentage of solvent B was increased as follows: from 2% to 4% in 4 min, to 8% in 2 min, to 25% in 64 min, to 40% in 12 min, to 80% in 4 min, followed by re-equilibration back to 2% B in 4 min (for the phosphoproteome). For the full proteome analysis, the steps were as follows: from 2% to 8% in 4 min, to 28% in 104 min, to 40% in 4 min, to 80% in 4 min, followed by re-equilibration back to 2% B in 4 min. The peptides were introduced into the Fusion Lumos via a Pico-Tip Emitter 360 µm OD x 20 µm ID; 10 µm tip (New Objective) and an applied spray voltage of 2.4 kV. The capillary temperature was set at 275°C. Full mass scan was acquired with mass range 375 to1400 m/z for the phosphoproteome (375 to 1500 m/z for the full proteome), in profile mode in the orbitrap with resolution of 120000. The filling time was set at maximum of 50 ms for the full proteome with a limitation of 4×105 ions. Data dependent acquisition (DDA) was performed with the resolution of the Orbitrap set to 30000, with a fill time of 110 ms for the phosphoproteome (94 ms for the full proteome) and a limitation of 1×105 ions. A normalized collision energy of 34 was applied. MS2 data was acquired in profile mode. Fixed first mass was set to 110 m/z.

### 4.2 Data analysis

If not stated otherwise statistical analysis was performed using the R programming language (version 4.4.1) in the RStudio environment (version 2024.04.2+764).

#### Image Analysis of Wide-Field Microscopy Images

Image inspection and analysis, including nuclei segmentation and fluorescence quantification, were performed using Fiji ImageJ (Schindelin et al. 2012) and CellProfiler (Stirling et al. 2021). Details of the CellProfiler pipeline and parameters are provided in the Github repository (CellProfiler Pipeline). Key steps included image loading, channel assignment, nuclei segmentation, and fluorescence measurement. Data was exported to CSV files for downstream analysis.

CellProfiler output was imported into R (R Foundation for Statistical Computing 2021) for further analysis. Treatments were assigned based on well and position information extracted from the filenames. Fluorescence intensity values were rescaled and background subtracted. Autofluorescence and non-fibrillar ECM staining corrections were applied. Intensity values were then normalized to the number of nuclei in each image, calculating a per-nucleus intensity value. Images with low nuclei counts were excluded from the analysis.

#### COL1 deposition analysis

A linear mixed model with random intercept was applied to analyze COL1 deposition over time, accounting for treatment (TGF-β or control), time (0, 12, 24, 48, 72, and 96 h), and their interaction as fixed effects, with plate (biological replicate) set as random effect. The model formula in R notation is:

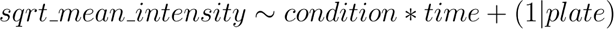

The 0 h time point values were duplicated to account for the absence of a 0 h TGF-β treatment. The model includes:

● An intercept (control condition at 0 h, varying per plate)
● A conditionTGF parameter (TGF-β treatment effect)
● Time parameters for each time point (time12h, time24h, etc.)
● Interaction terms between time and condition (conditionTGF:time12h, conditionTGF:time24h, etc.)

The treatment effect at any given time point is the sum of the relevant parameters (e.g., for TGF-β at 12 h: intercept + conditionTGF + time12h + conditionTGF:time12h).

Average COL1 intensity per cell was calculated for each condition, technical, and biological replicate. Square root transformation was applied to stabilize the variance and improve residual distribution (Piepho 2009). Analysis of variance using type III sum of squares, followed by a post hoc test, was performed to identify significant differences between factor levels. For visualization, sqrt_mean_int data was normalized per plate by subtracting the 0 h time point values. The resulting data points and model-predicted values were plotted (Figure 1C). The underlying data can be found in Table S8.

#### Analysis of ECM Deposition of siRNA validation experiments

For siRNA treatments, ECM staining per cell was normalized to the siNeg9 (non-targeting siRNA) control:

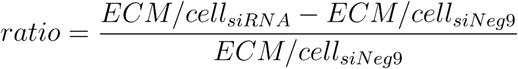

This normalization was performed separately for control and TGF-β treated samples, per plate (biological replicate). For TGF-β treated samples, an additional ratio was calculated by dividing siRNA + TGF-β treated samples by siNeg9 + TGF-β samples. Statistical significance was determined using unpaired two-sided t-tests. The underlying data can be found in Table S9.

#### RT-qPCR data statistical testing

For RT-qPCR analysis in the validation experiments (Results section 2.4), data preprocessing involved removing technical replicates with standard deviation > 0.5 if clearly outliers. The 2^-ΔΔCT^ method (Livak & Schmittgen 2001) was applied for analysis. CT values were normalized to GAPDH as the housekeeping gene, and ΔΔCT values were calculated relative to the siNeg9 control sample at 48 h post siRNA transfection. mRNA expression was presented as percentages, with the control set to 100%. Statistical analysis included unpaired two sided t-tests comparing 72 h and 96 h samples (siRNA control vs. siNeg9 control, and siRNA TGF-β vs. siNeg9 TGF-β). It should be noted that more technical replicates of siNeg9 samples were included as reference samples across different RT-qPCR plates, resulting in a higher number of data points for these controls in the analysis. The underlying data can be found in Table S10.

#### RNAseq alignment and preprocessing

Sequencing reads were aligned using STAR (version 2.7.9a) with default parameters on GRCh38. The gene count tables were produced during the alignment (--quantMode GeneCounts) using the annotation GRCh38.93. One outlier sample (ctrl, 24h, replicate A) was removed and one sample swap was corrected (24h and 48h TGF-β treated, replicate B) because of experimental errors. Lowly expressed genes were removed using the filterByExpr() function of the edgeR R package. Normalization factor calculation and normalization were performed using the calcNormFactors() and cpm() functions of the edgeR R package.

#### MS database search

For the secretomics data, all raw files were converted to mzML format using MSConvert from Proteowizard (version 3.0.22129), using peak picking from the vendor algorithm. Files were then searched using MSFragger v3.7 (Kong et al. 2017) in Fragpipe v19.1 against the Swissprot Homo sapiens database (20,594 entries) containing common contaminants and reversed sequences. The standard settings of the Fragpipe TMT11 workflow were used. The following modifications were included into the search parameters: Carbamidomethyl (C) and TMT11 (K) (fixed modification), Acetyl (Protein N-term), Oxidation (M) and TMT11 (N-term) (variable modifications). For the full scan (MS1) a mass error tolerance of 10 ppm and for MS/MS (MS2) spectra of 0.02 Da was set. Further parameters were set: Trypsin as protease with an allowance of maximum two missed cleavages and a minimum peptide length of seven amino acids was required. The false discovery rate on peptide and protein level was set to 0.01.

For the phosphoproteomics data, MSFragger v3.8 (Kong et al. 2017) was used to process the acquired data, which was searched against the homo sapiens Uniprot proteome database (UP000005640, ID9606, 20594 entries, release October 2022) with common contaminants and reversed sequences included. The following modifications were considered as fixed modification: Carbamidomethyl (C) and TMT16 (K). As variable modifications: Acetyl (Protein N-term), Oxidation (M) and TMT16 (N-term), for the phosphoproteome specifically phosphorylation on STY. For the MS1 and MS2 scans a mass error tolerance of 20 ppm was set. Further parameters were: Trypsin as protease with an allowance of maximum two missed cleavages; Minimum peptide length of seven amino acids; at least two unique peptides were required for a protein identification. The false discovery rate on peptide and protein level was set to 0.01.

#### Proteomics preprocessing

For the secretomics data, the raw output files of MSFragger (Kong et al. 2017) (protein.tsv – files) were processed using the R programming language (R Foundation for Statistical Computing 2021). Contaminants including albumin were filtered out and only proteins that were quantified with at least two unique peptides were considered for the analysis. Moreover, only proteins which were identified in two out of three mass spec runs were kept. 2188 proteins passed the quality control filters.

For the phosphoproteomics data, the raw output files of MSFragger (Kong et al. 2017) (psm.tsv for phospho data and protein.tsv files for input data) were processed using the R programming language. Only peptide spectral matches (PSMs) with a phosphorylation probability greater 0.75 and proteins with at least 2 unique peptides were considered for the analysis. Phosphorylated amino acids were marked with a * in the amino acid sequences behind the phosphorylated amino acid, labeled with a 1, 2 or 3 for the number of phosphorylation sites in the peptide and concatenated with the protein ID in order to create a unique ID for each phosphopeptide. Raw TMT reporter ion intensities were summed for all PSMs with the same phosphopeptide ID. For the input data, the reporter ion intensities were used as given in the protein.tsv output files. Phospho signals were also normalized by input abundance. For this the reporter ion intensity for each unique phospho ID, condition and replicate was normalized according to the following formula:

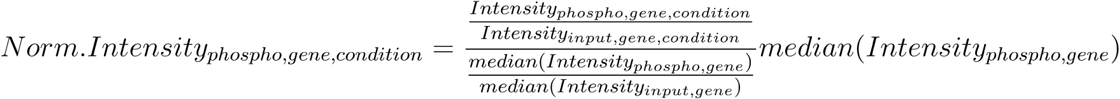

For all proteomics data modalities, transformed summed TMT reporter ion intensities or log2 transformed raw TMT reporter ion intensities were first cleaned for batch effects using the ‘removeBatchEffects’ function of the limma package (Ritchie et al. 2015) and further normalized using the vsn package (variance stabilization normalization - (Huber et al. 2002)). For observations with less than 30% missing values in the phosphoproteomics data, missing values were imputed with the ‘knn’ method using the Msnbase package (Gatto & Lilley 2012).

#### Differential abundance analysis

Proteins and transcripts were tested for differential expression using the limma package. For the proteomics data modalities, phospho, normalized phospho and input data was tested separately. For the secretomics data, the replicate information was added as a factor in the design matrix given as an argument to the ‘lmFit’ function of limma. Resulting p-values were adjusted for multiple testing. A protein was considered statistically significantly different with an adjusted p-value below 0.05 and log2 fold change above log2(1.5) or below log2(1/1.5). A transcript was considered statistically significantly different with an adjusted p-value below 0.05 and log2 fold change above log2(2) or below log2(2). For the secretomics data, the proteins were filtered for secreted proteins using the gene sets “M5889” and “M5885” from the MSIGDB (Bhuva et al. 2024).

#### Correlation analyses

Technical (Pearson) correlation between all samples and replicates was computed using the counts and reporter intensities before differential expression analysis. To correlate significantly affected molecules between data sets and conditions, the t-values of proteins that were affected in at least one condition were correlated using Pearson correlation, with the exception of the phosphoproteomics data for which this has been done separately.

#### Pathway enrichment analysis

For the path enrichment analysis, MSIGDB (Bhuva et al. 2024) Hallmark pathways were used with the decoupleR package (Badia-I-Mompel et al. 2022) (normalized weighted average method) to calculate path enrichment values from log2 fold change values.

#### Enzyme activity analysis

In order to estimate the activity of different input nodes (transcription factors (TFs), kinases, phosphatases) the normalized weighted mean and mixed linear model (mlm) methods of the decoupleR package were used. For transcription factors, the DOROTHEA database was used to obtain TF-target interactions with a confidence level of A, B or C (Garcia-Alonso et al. 2019). The interactions were downloaded using the OmnipathR package (Türei et al. 2016). Phosphosite-enzyme collections were likewise downloaded with OmnipathR. For the decoupleR tool, log2 fold-change values of transcripts and phosphosites after limma analysis were used as input per condition. A TF, kinase or phosphatase was considered to be significantly affected for p-value < 0.03 and absolute normalized enrichment score > 3.

#### Network modeling

The primary goal of the network modeling was to extract a subnetwork that could consistently connect nodes from different data modalities in both sign and direction, using a template PKN as starting point using COSMOS (Dugourd et al. 2021). The PKN contains the signed and directed protein-protein interactions (node A activates/inactivates node B) which are used to represent signaling interactions between root nodes (input nodes) and downstream nodes (measurement nodes) The subnetwork is selected through an integer linear programming (ILP) formulation that solves a particular instance of the Prize-Collecting steiner tree problem with multiple root nodes and additional custom constraints (see GitHub Repo).

In this study, we used a prior knowledge network which is provided as part of the OmnipathR package using the import_all_interactions() function. This initial PKN was filtered for trusted resources and interactions with a valid consensus signal. Additionally, the core of the TGFb signaling module was fixed to guide the optimisation (enforce TGFb to SMAD1-5, support TGFb to MAPK1, MAPK14, AKT1 and PI3K) following Park and Yoo (Park & Yoo 2022). The final PKN (42k interactions) was obtained by filtering for expression, downstream neighbors and consistent TF-target interactions as reported before (Dugourd et al. 2021).

As input we used significant transcription factors and kinases/phosphatase from the activity estimation analysis which we divided into early and late groups (0.08 h, 1 h, 12 h and 24 h, 48 h, 72 h, 96 h) as well as secretomics hit filtered for proteins which are known to be secreted (0.08 h, 1 h, 12 h, 24 h for early network, 48 h, 72 h, 96 h for late network). This list was manually extended with COL1A1, COL5A1, SERPINE2, SPARC, ITGB1, VIM, JUP, ACTA1, HSPG2, LOXL2, TNC, TGFBI, IGFBP3, IGFBP7, LTBP2, TAGLN, CCN2, LRATD2, MRC2, FN1, FBN1, BGN, ADGRG1, MMP2, ITGA11, AMIGO2, ADAM12, CPA4, DCDC2, PLEKHG4 and ITGB1BP1 which were significantly deregulated in the secretomics experiment and reported to be secreted before. The number of input nodes has been chosen to find an acceptable compromise of computational cost and information content for the network modeling step.

In total, four subnetworks were inferred based on different time points and input modalities to account for the time-resolved nature of the obtained data. To start the optimisation we used the stimulating agent TGFb as upstream input node of early affected enzymes. This network was combined with a second network connecting the set of early affected enzymes and early secretomics hits. This combination represents early signaling events stimulated by TGFb treatment which then lead to altered protein secretion up to 24 hours (early network). To model assumed autocrine function of secreted factors we modeled later signaling processes by inferring a first network between secreted proteins included in the early network and enzymes affected at a later time point (between 24 h and 96 h). In a last step this network was combined with a fourth subnetwork connecting later affected enzymes with late secretomics hits (late network).

#### Network level analyses

Node enrichment analysis for the early and late network was performed using ReactomePA (Gillespie et al. 2022). The enrichment results were compared by calculating the log2 odds ratio for each pathway. Closeness centrality was computed per subnetwork using the closeness centrality method implemented in the igraph R-package (http://igraph.org).

## Acknowledgements

We would like to acknowledge the invaluable contributions of several core facilities and individuals who made this research possible:

The EMBL Genome Core Facility, particularly Vladimir Benes, Jonathan Landry, and Ferris Jung, for their expertise in transcriptomics. We thank the EMBL Advanced Light Microscopy Facility, notably Aliaksandr Halavatyi, Beate Neumann, and Manuel Gunkel, for their assistance with imaging. For statistical analysis, we are grateful to Lukas Josef Koppensteiner from BOKU Vienna, and Charlotte Boys, Robin Fallegger and Rebecca Terrall Levinson from the Saez lab. Furthermore, we want to thank Isabel Kemmer from the Euro-BioImaging ERIC for support in bioimage data upload.

We appreciate Kim Remans from the EMBL Protein Expression and Purification Core Facility for providing the protocol for TCA precipitation (secretomics) and fluorescent dye (CNA35).

We also extend our thanks to Christoph A. Merten from the Institute of Bioengineering, School of Engineering, École Polytechnique Fédérale de Lausanne (EPFL), Lausanne, Switzerland, for his contributions.

The collaborative efforts of these individuals and institutions were instrumental in the success of this research. Their expertise, guidance, and support have significantly enhanced the quality and scope of our work.

## 5. Conflict of interests

JSR reports funding from GSK, Pfizer and Sanofi and fees/honoraria from Travere Therapeutics, Stadapharm, Astex, Pfizer, Moderna, Grunenthal and Owkin.

## 6. Authors contributions

Nadine Tuechler: Investigation, Conceptualization, Validation, Formal analysis, Visualization, Project administration, Methodology, Writing - Original Draft Preparation Mira Lea Burtscher: Software, Formal analysis, Visualization, Methodology, Writing - Original Draft Preparation

Martin Garrido-Rodriguez: Formal analysis, Visualization, Writing - Review & Editing

Muzamil Majid Khan: Conceptualization, Writing - Review & Editing

Denes Türei: Conceptualization, Writing - Review & Editing

Christian Tischer: Ressources, Methodology, Visualization

Sarah Kaspar: Software, Formal analysis, Methodology

Jennifer Jasmin Schwarz: Methodology, Resources

Frank Stein: Methodology, Software, Ressources

Mandy Rettel: Methodology, Ressources

Rafael Kramann: Ressources, Writing - Review & Editing

Mikhail Savitski: Supervision, Writing - Review & Editing

Julio Saez-Rodriguez: Supervision, Project administration, Funding acquisition, Writing - Review & Editing

Rainer Pepperkok: Supervision, Ressources, Project administration, Funding acquisition, Writing - Review & Editing

## 7. Data and code availability

The mass spectrometry proteomics data have been deposited to the ProteomeXchange Consortium via the PRIDE partner repository with the dataset identifier PXD056096. The transcriptomics sequencing data has been deposited to the ArrayExpress repository with the ArrayExpress accession E-MTAB-14521. Microscopy data are available in the BioStudies database (https://www.ebi.ac.uk/biostudies/) under accession number S-BIAD1415 (DOI 10.6019/S-BIAD1415).

All code used to perform the computational analyses described and to reproduce the figures is available at https://github.com/saezlab/kidneyfibrosis_multiomicsmodel_paper. All figures and analyses can be reproduced using the provided supplementary tables (contain processed data and analysis results) and the source data provided on Github.

## 8. Supplementary Materials

**Supplementary Figure 1.**
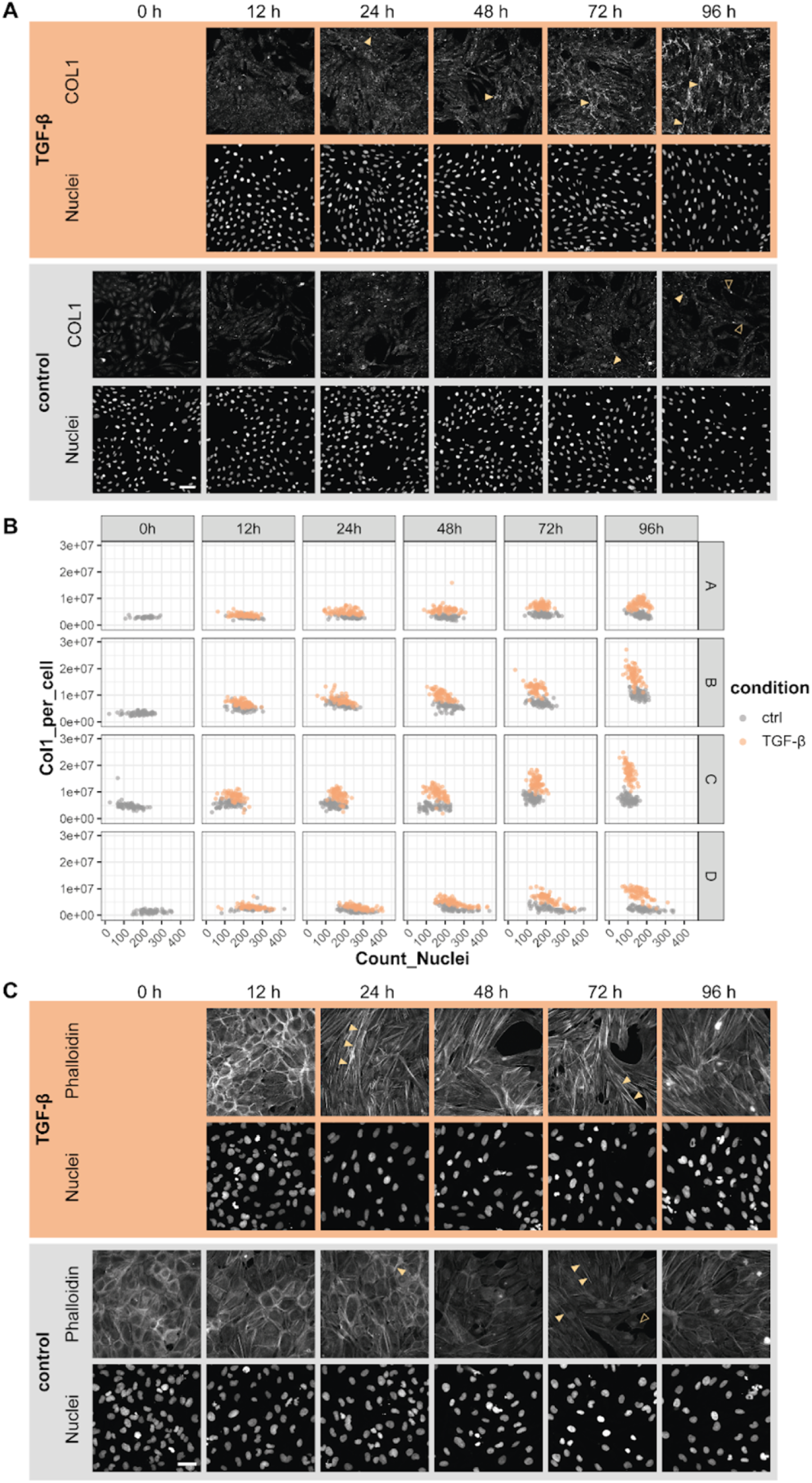
(A) Widefield microscopy images showing COL1 (top rows) and nuclear (Hoechst, bottom rows) staining in control and TGF-β-treated conditions at 0, 12, 24, 48, 72, and 96 hours. (B) Quantification of COL1 expression per cell over time. Cells were cultured in four biological replicates (A-D) ± TGF-β for 0-96 h. Data points represent individual images, color coded for control (grey) and TGF-β (orange) conditions. Y-axis shows normalized COL1 intensity per cell, x-axis shows nuclei count per image. Images with less than 20 nuclei were excluded from the analysis. (C) Sum Z-projections of confocal microscopy images displaying F-actin (phalloidin, top rows) and nuclear (bottom rows) staining in control and TGF-β treated conditions at the same time points as in (A). In panels (A) and (C), the control condition is shown in the lower half (gray background), while the TGF-β treated condition is presented in the upper half (orange background). Scale bar = 100 µm. Yellow arrowheads indicate specific features of interest (fibrillar collagen or F-actin, respectively), while hollow arrowheads indicate autofluorescence of cells.

**Supplementary Figure 2.**
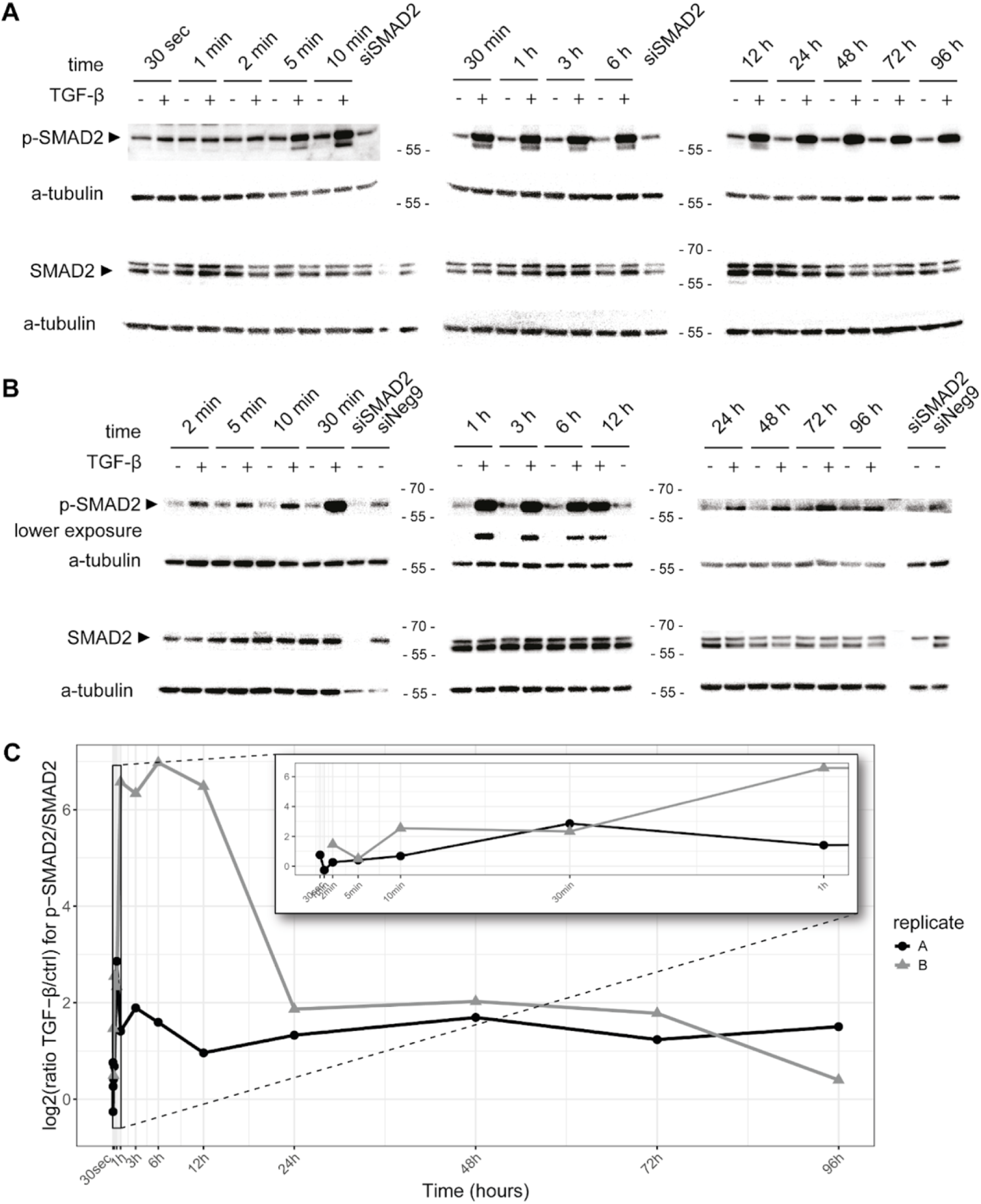
(A) Western blot analysis of phosphorylated SMAD2 (p-SMAD2), total SMAD2, and α-tubulin (loading control) in response to TGF-β treatment over time. Time points range from 30 seconds to 96 hours. siSMAD2 condition is included as a control. (B) Western blot analysis of a biological replicate, similar to (A). Time points range from 2 minutes to 96 hours. siSMAD2 and siNeg9 conditions are included as controls. (C) Quantification of p-SMAD2 levels normalized to total SMAD2 from (A) and (B). Each of the measurements was normalized to the corresponding loading control before calculating the log2 fold change between TGF-β and control treated samples. The graph shows the log2 ratio of TGF-β/ctrl for p-SMAD2/SMAD2 across all time points. Two replicates (A and B) are represented.

**Supplementary Figure 3.**
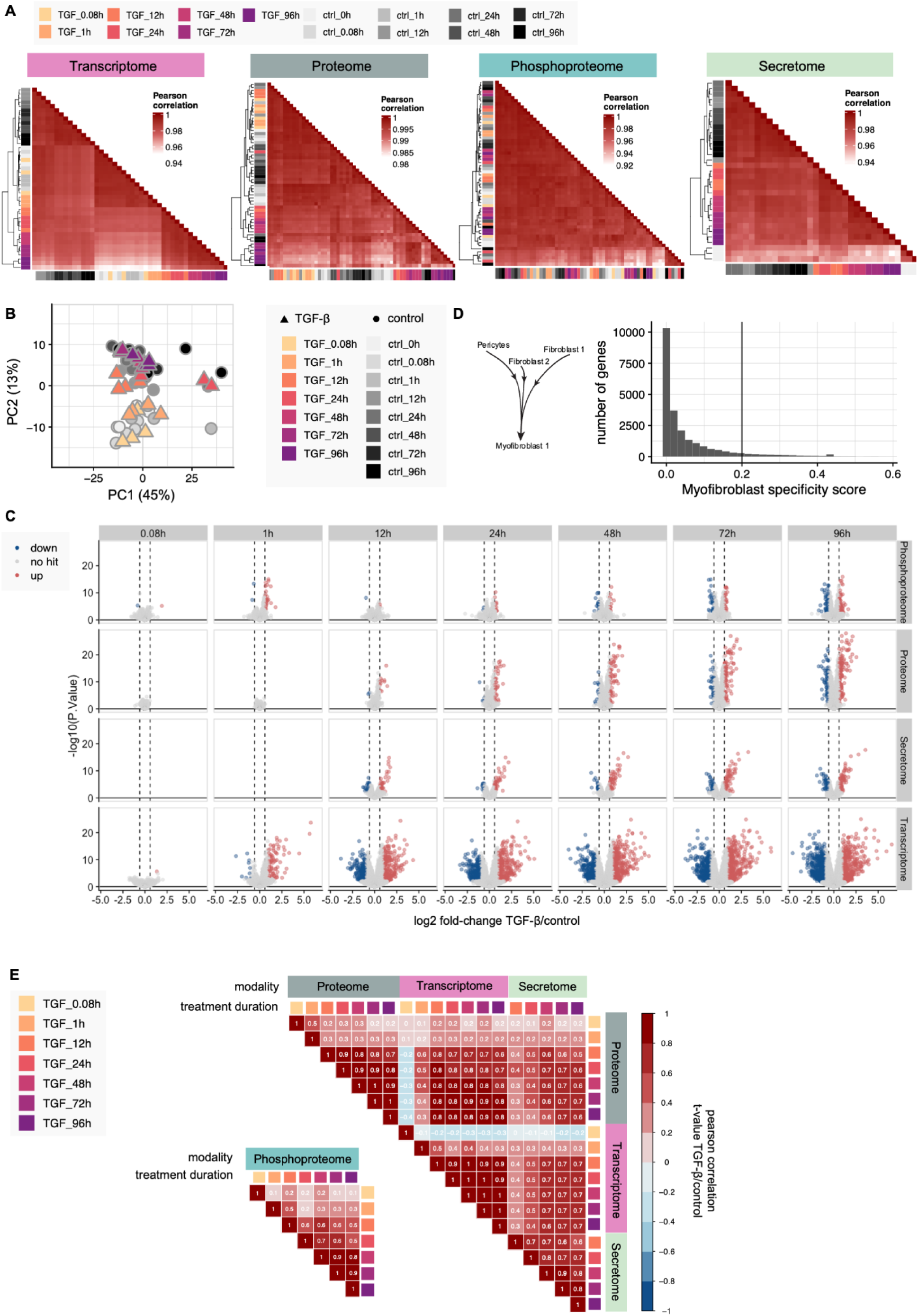
(A) Heatmap of Pearson correlation of TMT reporter intensities or gene counts for each omics modality across all samples and replicates. The color gradients indicate the time points for TGF-β-treated (yellow to purple) and control (greyscale) samples. (B) PCA scatter plot for phosphoproteomics data (PC1 vs PC2). Triangles represent TGF-β-treated samples, circles represent controls. Color gradients indicate time points as in (A). (C) Results of the differential expression analysis per time point and omics modality. Colors indicate transcripts and proteins significantly deregulated in abundance upon TGF-β stimulation in comparison to control samples (limma, adjusted p-value < 0.05, absolute log2 fold-change > log2(2) for transcripts, absolute log2 fold-change > log2(1.5) for proteins). (D) Specificity score distribution for myofibroblasts from CKD patients (Human PDGFRβ+ level 2) retrieved from Kuppe et al. 2021 (Kuppe et al. 2021). The black line indicates the chosen specificity cutoff for comparisons to this study (0.2). (E) Heatmap showing Pearson correlations between time points and omics modalities for genes with a significantly deregulated transcript or protein in at least one modality, filtered for genes detected in all modalities (limma, adjusted p-value < 0.05, absolute log2 fold-change > log2(2) for transcripts, absolute log2 fold-change > log2(1.5) for proteins).

**Supplementary Figure 4.**
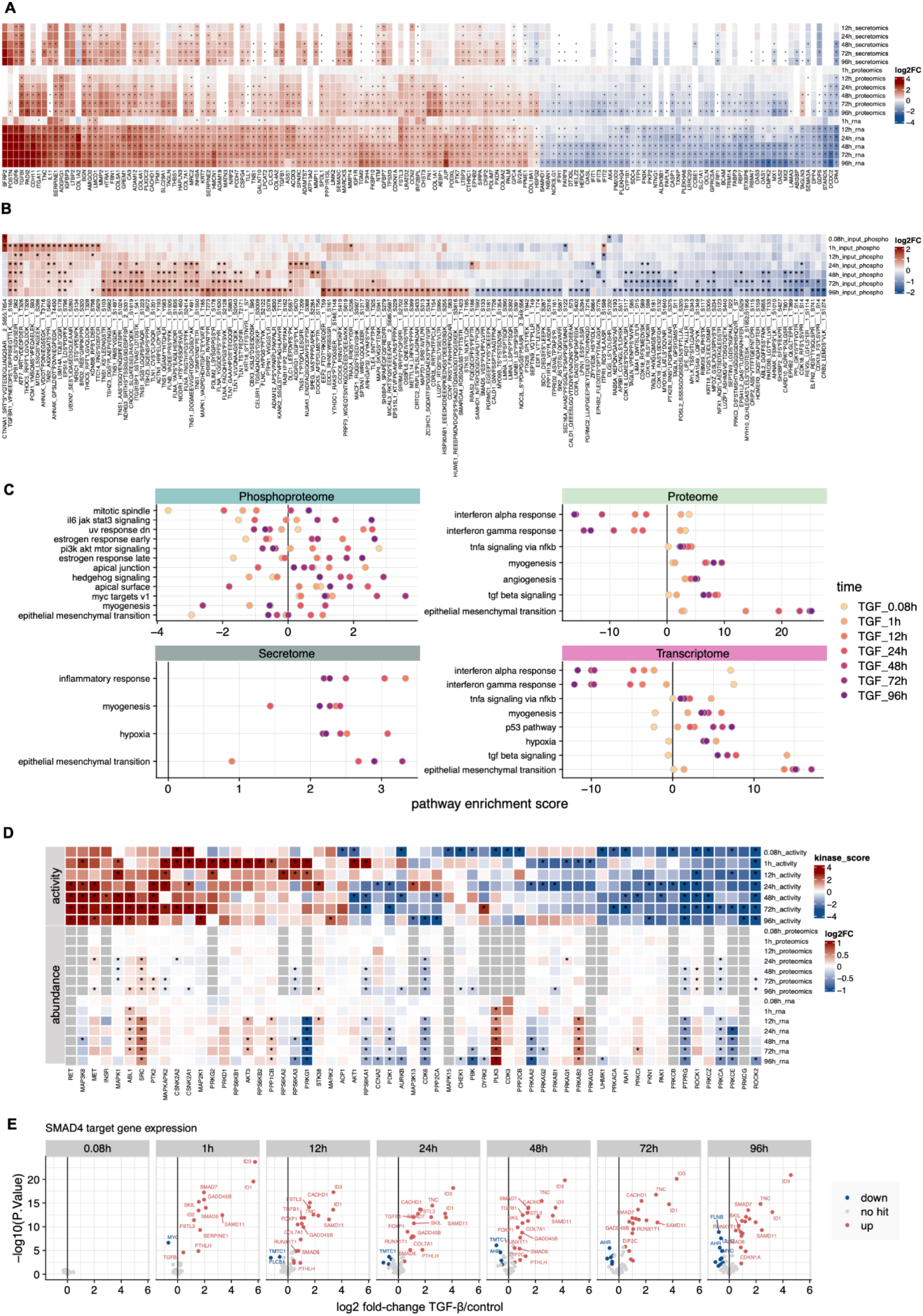
(A) Heatmap of differential abundance per time point of transcripts, proteins and secreted proteins across time points. Included are genes significantly affected in at least two modalities upon TGF-β stimulation (limma, adjusted p-value < 0.05, absolute log2 fold-change > log2(2) for transcripts, absolute log2 fold-change > log2(1.5) for proteins). The color intensity indicates magnitude and direction of change. (B) Heatmap displaying differential abundance per time point of the top affected phosphopeptides upon TGF-β stimulation. The color indicates the direction of change. (C) Dot plots of top significantly enriched pathways (GSEA using decoupleR with MSIGDB reactome database, p-value < 0.05) per time point and omics modality. Color represents the time points. (D) Heatmap of differential protein and transcript abundance with matched activity scores of kinases/phosphatases per time point. Kinases/phosphatases were considered if they showed significantly altered activity upon TGF-β stimulation (enzyme activity enrichment analysis using decoupleR, p-value < 0.05 and absolute enrichment score > 3). (E) Exemplary target profile of the transcription factor SMAD4 over time. Each point represents a known SMAD4 target transcript, with color indicating up- or down-regulation. The y-axis shows statistical significance. The differential abundance signal of these known target transcripts is summarized to an activity score per time point.

**Supplementary Figure 5.**
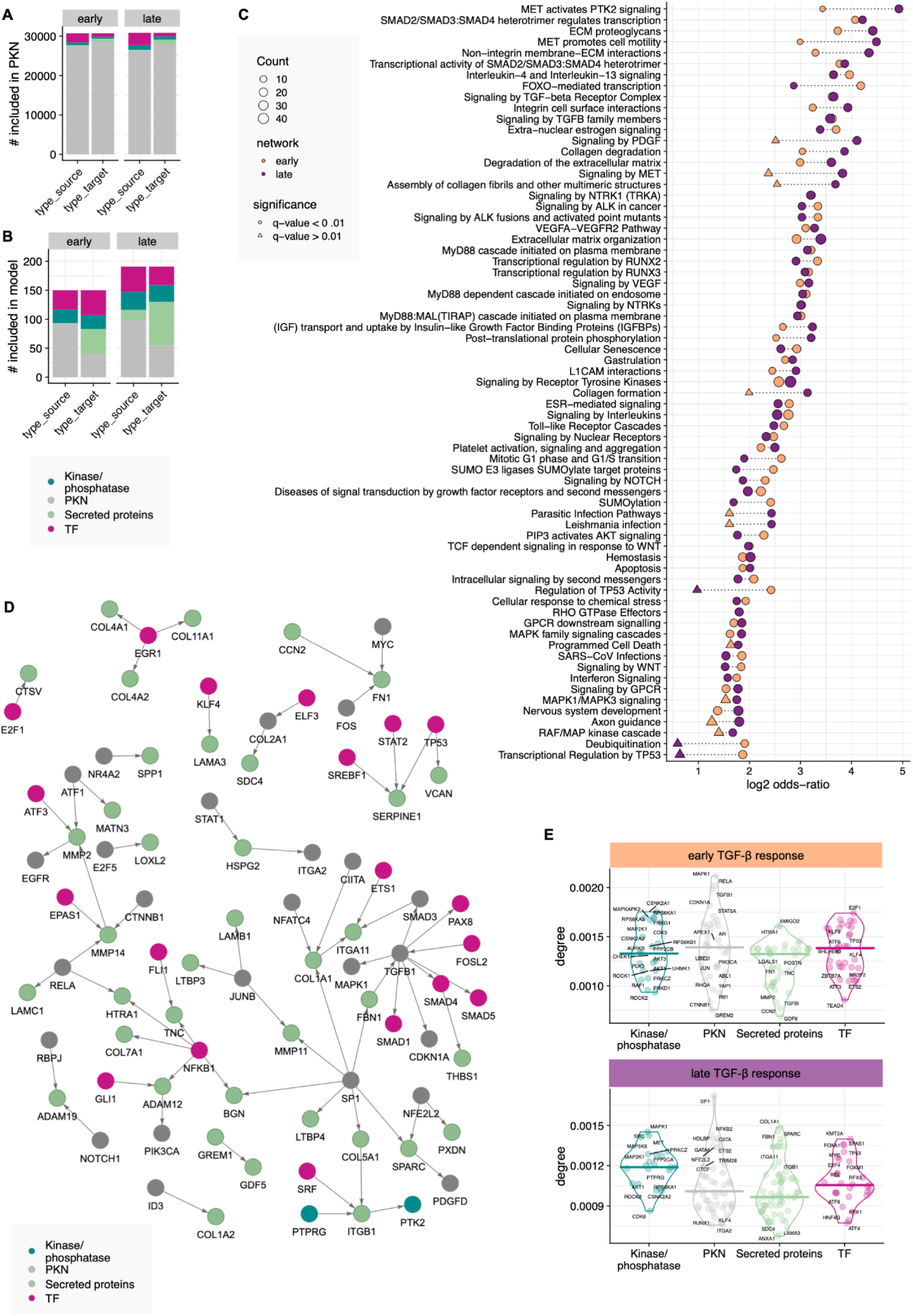
(A) Overview of node types in the prior knowledge network. All modalities are reflected as protein-protein interaction source and target for the early and late TGF-β response model. B) Overview of node types in the obtained solution network models for early and late TGF-β response. The node type for protein-protein interaction source and target nodes reflect the chosen hierarchy. C) Significant pathways in early and late TGF-β response network model (pathway overrepresentation analysis of network nodes using the Reactome pathway database, p-value < 0.05). Point color indicates the network for which the enrichment has been calculated. Point size shows the number of nodes in the network mapped to the pathway. D) Reactome TGF-β signaling pathway in network models shows links between different node types. Node color indicates node type. E) Members of the Reactome TGF-β signaling pathway in the differential abundance data upon TGF-β stimulation. Color indicates direction of change.

**Supplementary Figure 6.**
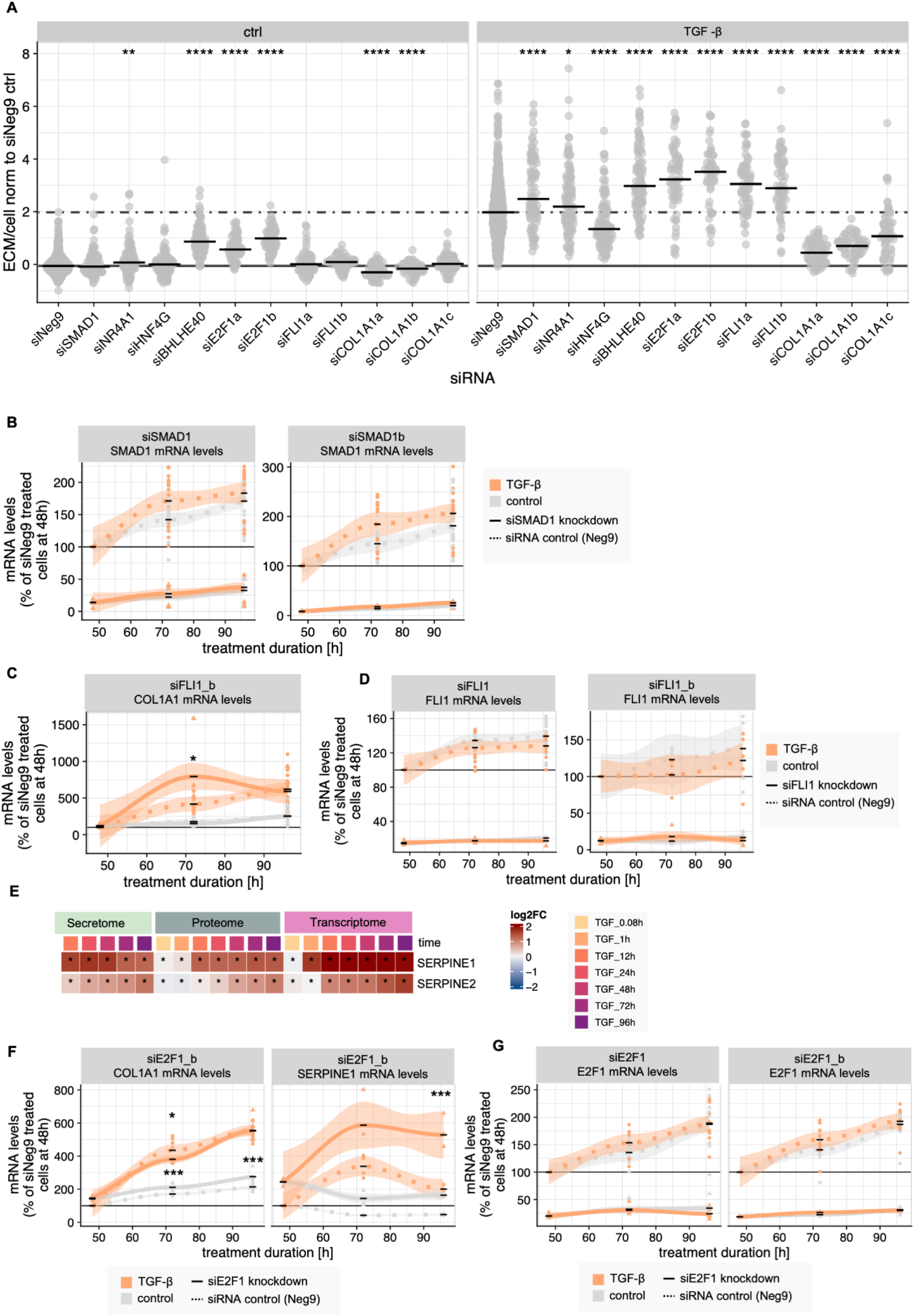
(A) Fluorescence intensity of ECM for both untreated (ctrl) and TGF-β stimulated conditions for all performed knockdowns (96h knockdown followed by 48h +/- TGF-β treatment). Intensities of each knockdown were compared to the siNeg9 control condition (+/- TGF-β treatment) using a t-test (* corresponds to p-value < 0.05, ** corresponds to p-value < 0.01, *** corresponds to p-value < 0.001). For panels B-D and F, color indicates TGF-β stimulation (orange) vs control (gray). Line type distinguishes between siNeg9 control (solid) and target gene knockdown (dashed). (B) RT-qPCR data showing SMAD1 knockdown efficiency for various mRNAs across different time points. (C) RT-qPCR results demonstrating the effect of FLI1 knockdown (using a second set of siRNA) on its potential downstream target COL1A1 at different time points. (D) RT-qPCR data confirming FLI1 knockdown efficiency for multiple mRNAs at various time points. (E) Differential expression analysis results for SERPINE1 and SERPINE2 in the secretomics, proteomics and transcriptomics data per time point. Significance is indicated by one star (limma, adjusted p-value < 0.05, absolute log2 fold-change > log2(1.5) or absolute log2 fold-change > log2(2) for transcriptomics data). (F) RT-qPCR data to confirm E2F1 knockdown effect on its potential downstream targets COL1A1 and SERPINE1 +/-TGF-β stimulation at different time points using a second set of siRNA.(G) RT-qPCR data to confirm E2F1 knockdown efficiency for different mRNAs (columns) +/-TGF-β stimulation at different time points.

Table S1 - differential abundance analysis results

Table S2 - pathway enrichment results

Table S3 - TF and kinase enrichment results

Table S4 - network models edge table

Table S5 - network models node table

Table S6 - qPCR primer sequences

Table S7 - siRNA sequences

Table S8 - COL1 imaging data

Table S9 - COL1 + siRNA imaging data

Table S10 - qPCR results

All supplementary tables can be found in the provided excel table Supplementary_Data.xlsx

## Notes

https://github.com/saezlab/kidneyfibrosis_multiomicsmodel_paper

